# Functional modeling of NMIHBA-causing *PRUNE1* variants reveals a requirement for its exopolyphosphatase activity

**DOI:** 10.1101/2020.03.02.973909

**Authors:** Harikiran Nistala, John Dronzek, Claudia Gonzaga-Jauregui, Shek Man Chim, Saathyaki Rajamani, Samer Nuwayhid, Dennis Delgado, Elizabeth Burke, Ender Karaca, Matthew C. Franklin, Prasad Sarangapani, Michael Podgorski, Yajun Tang, Melissa G. Dominguez, Marjorie Withers, Ron A. Deckelbaum, Christopher J. Scheonherr, William A. Gahl, May C. Malicdan, Brian Zambrowicz, Nicholas W. Gale, Richard A. Gibbs, Wendy K. Chung, James R. Lupski, Aris N. Economides

## Abstract

Neurodevelopmental disorder with microcephaly, hypotonia, and variable brain anomalies (NMIHBA) is an autosomal recessive neurodevelopmental and neurodegenerative disorder characterized by global developmental delay and severe intellectual disability. Microcephaly, progressive cortical atrophy, cerebellar hypoplasia and delayed myelination are neurological hallmarks in affected individuals. NMIHBA is caused by biallelic variants in *PRUNE1* encoding prune exopolyphosphatase 1. We provide in-depth clinical description of two affected siblings harboring compound heterozygous variant alleles, c.383G>A (p.Arg128Gln), c.520G>T (p.Gly174*) in *PRUNE1*. To gain insights into disease biology, we biochemically characterized missense variants within the conserved N-terminal aspartic acid-histidine-histidine (DHH) motif and provide evidence that they result in the destabilization of protein structure and/or loss of exopolyphosphatase activity. Genetic ablation of *Prune1* results in midgestational lethality in mice, associated with perturbations to embryonic growth and vascular development. Our findings suggest that NMIHBA results from hypomorphic variant alleles in humans and underscore the potential key role of PRUNE1 exopolyphoshatase activity in neurodevelopment.

## Introduction

Prune exopolyphosphatase 1 (PRUNE1) belongs to the DHH (Asp-His-His) superfamily of proteins, which includes phosphoesterases, exopolyphosphatases and pyrophosphatases (1-3). Members of this superfamily are characterized by the DHH motif within the N-terminal domain that contains highly conserved charged residues required for binding metal cofactors and enzymatic function (4). PRUNE1 is ubiquitously expressed across multiple tissues with widespread central nervous system expression during fetal development in humans (5). Previous studies have implicated PRUNE1 in a complex network of interactions regulating cell cycle and motility, with elevated *PRUNE1* expression associated with cancer metastasis (6-8).

Karaca et al., reported five individuals from four unrelated families segregating biallelic predicted pathogenic variants in *PRUNE1* with a novel neurodevelopmental disorder characterized by microcephaly, hypotonia, and variable brain anomalies (NMIHBA, MIM #617481)(9). The authors reported two different homozygous missense variants (p.D30N and p.D106N) in three consanguineous families and compound heterozygous (p.R128Q; p.G174*) variants in a non-consanguineous European ancestry family. Several groups have since identified other disease associated recessive alleles (p.P54T; p.R297W; c.521-2A>G: IVS4-2A>G) clustering throughout the conserved N-terminal domain. The observed incidence of NMIHBA in geographically distant families establishes this disorder as a pan-ethnic neurodevelopmental disorder (8, 10, 11). In addition to its nondescript developmental delay (DD)/intellectual disability (ID) nervous system hallmarks, NMIHBA patients may manifest more variable clinical features including craniofacial anomalies (sloped forehead, large prominent ears, prominent eyes and narrow palate), scoliosis, joint contractures, muscle atrophy, hyperreflexia, profound global developmental delay, fronto-temporal atrophy, cortical and cerebellar atrophy, seizures, and spastic quadriplegia.

Although the neurodevelopmental and neurobiological etiology of NMIHBA remains unclear, the severe clinical and developmental aspects of this disorder suggest an important role for PRUNE1 in fundamental cellular functions. Indeed, prior studies have implicated PRUNE1 in cell proliferation and migration, processes mediated in part through interactions with nucleoside diphosphate kinase 1 (NME1), glycogen synthase kinase-3β (GSK-3β) and β-tubulin respectively (4, 8, 12). Although *in silico* predictions of likely damaging effects of missense alleles classify them as likely pathogenic, the direction of effect (i.e. loss-of-function versus gain-of-function) and functional consequences of these variants have not been established. The observation of biallelic variants for a recessive disease trait, and the identification of putative loss of function splicing and nonsense variants support the notion of a reduced or loss of PRUNE1 function in NMIHBA pathophysiology. In agreement, patients harboring compound heterozygous mutations p.R128Q and p.G174* (described herein) manifest typical NMIHBA similar to those patients with homozygous missense variants spanning the conserved DHH (p.D30N, p.P54T, p.D106N, p.R128Q) and DHHA2 domains (p.R297W) (8-10). Heterozygous carriers of reported pathogenic missense and truncating variants in *PRUNE1* do not appear to have any phenotypic manifestations in the spectrum of NMIHBA, indicating that *PRUNE1* haploinsufficiency in heterozygous carrier parents is permissive for neurotypical development. To gain further neurodevelopmental and neurobiological insights into NMIHBA, we investigated the functional effects of *PRUNE1* variant alleles and generated a knock-out mouse model of *Prune1*.

We undertook comprehensive clinical phenotyping of two siblings from one of the initially reported families (SZ51) with NMIHBA and summarize the phenotypic spectrum across all NMIHBA patients reported to date. Additionally, we provide experimental evidence that disease associated missense variant alleles that map within the conserved N-terminal domain can result in destabilization of the protein structure and consequent loss of exopolyphosphatase activity. Furthermore, we demonstrate that homozygous null alleles in *Prune1*^*-/-*^ mice results in multiple vascular anomalies including abnormal vascular network within the yolk sac and disrupted cephalic vascular plexus culminating in midgestational lethality (E9.5). Although, Embryonic lethality of *Prune1* homozygous null mice precluded molecular exploration of the neurodevelopmental phenotypes, our study provides a potential biochemical basis to NMIHBA pathophysiology and implicitly an avenue for therapeutic exploration.

## Material and Methods

### Cell culture

HEK293 and human fibroblast culture medium consisted of DMEM (Gibco; 11995-065) supplemented with 10% fetal bovine serum (Gibco 16000-036), 1% non-essential amino acids (Gibco; 11140-050), and 1% Penicillin-Streptomycin (Gibco 15140-122). Cells were dissociated using 0.05% Trypsin-EDTA (Gibco 25300-054). HEK293 cells were seeded in 96 well plates at a density of 20,000 cells/well and in other formats (12 and 24-well) at a density of 50,000 cells/cm^2^. Fibroblasts were seeded at a density of 3000 cells/cm^2^, in all cases. Genotypes of patient-derived fibroblasts and control dermal fibroblasts (ATCC; PCS-201-010) were Sanger di-deoxy sequence verified using primers listed in supplementary Table S1.

### Immunoblotting and protein stability analyses

N-terminal HA-tagged PRUNE1 overexpression constructs were transfected into PRUNE1 non-expressing HEK293 cells using X-tremeGENE HP DNA (Roche; 06366236001) transfection reagent. As a transfection control CMV-driven eGFP expression construct was co-transfected at a 1:19 ratio. Two days later, whole cell lysate was collected, and total protein concentration was quantified using bicinchoninic acid assay. PRUNE1 levels in overexpressing HEK293 and in human fibroblasts were analyzed by immunoblotting using anti-PRUNE1 (Origene; TA344725) and/or anti-HA (Abcam; 18181) antibodies at 1:1000 dilution, using GAPDH (CST; 3683S) as a loading control.

Stability of the various PRUNE1 muteins was evaluated by overexpressing N-terminal HA-tagged PRUNE1 constructs in HEK293 cells. After 48 hours, medium containing either cycloheximide (100ug/mL; Sigma 1810) and/or MG132 (15uM; Sigma M8699) was added. Whole cell lysates were collected 0, 6 and 24 hours of cycloheximide treatment. Whole cell lysate from cells treated with both cycloheximide and MG132 were collected after 24 hours of treatment. Total protein concentration was quantified and immunoblotted using Li-COR Odyssey CLx imaging system. In complementary immunoblotting experiments, endogenous PRUNE1 levels in Sanger verified (Fig S2B) patient-derived fibroblasts BAB3500 (*PRUNE1*^*D106N/D106N*^) and UDP760 (*PRUNE1*^*R128Q/G174\**^) was compared to control neonatal dermal fibroblasts (ATCC, PCS-201-010).

### GloSensor cAMP and cGMP Assays

HEK293 cells were plated in poly-D-lysine coated 96 well plates (Corning 354651). The following day cells were co-transfected with 12.5ng pGloSensor-22F or pGloSensor-42F and 10ng of PRUNE1 overexpression constructs (D30N, D106N, R128Q, 4DD) or control PS100013, PDE4A or PDE5A vectors per well. Total DNA was maintained at 100ng per well using pUC19. Two days later, cells were changed into GloSensor equilibration medium (10% fetal bovine serum and 2% GloSensor assay reagent (Promega E1290) in CO_2_-independent medium (Gibco 18045-088). Upon equilibration at room temperature for 2 hours, basal luminescence was acquired using a PerkinElmer 2104 multimode plate reader. Forskolin (Sigma F3917) or sodium nitroprusside dihydrate (Millipore 567538) was added to the medium to a final concentration of 10µM or 50µM. cAMP or cGMP phosphodiesterase activity was evaluated by measuring luminescence recorded at 1-minute intervals up to 45 minutes or 60 minutes respectively.

### Exopolyphosphatase Assay

Exopolyphosphatase activity of PRUNE1 against short chain polyphosphate (PolyP) substrates, sodium tripolyphosphate (P3, Sigma 72061) and sodium tetraphosphate (P4, BOC Sciences 7727-67-5) was determined by a fixed time assay using BIOMOL GREEN phosphate detection kit (BML-AK111, Enzo Lifesciences). Reaction was performed 15-120s in 50µL of 100mM Tris/HCl, 50µM EGTA, 2mM MgCl_2_, pH 7.2 at substrate concentrations ranging from 0.5µM to 100µM and 1µM to 50µM for P3 and P4 respectively. Reactions were terminated by addition of 100uL of BIOMOL GREEN reagent, absorbance at 620nm was measured following a 30min incubation at 25°C. Kinetic parameters were assessed by plotting Lineweaver-Burk plots using GraphPad PRISM 7 software.

Exopolyphosphatase activity of wild-type PRUNE1 on medium and long chain PolyP substrates (consisting of 45, 65 and 700 orthophosphate monomers, Sigma S4379, Sigma S6253, Kerafast EUI002) was assessed by comparing to *S. cerevesia* PPX1 (Bioorbyt; ORB419100). 50nM of PRUNE1 or 100nM PPX1 were incubated with polyphosphates at a final concentration of 2mM (in terms of phosphate residues) in 100mM Tris-HCl, pH 7.2, 50µM EGTA, 2mM MgCl_2_ or 50mM HEPES/KOH, pH 8.0, 1mM MgCl_2_, 125 mM KCl respectively. Reactions were allowed to proceed at 37°C for 150 min. Reactions were terminated at 15, 30, 60 and 150min. Negative control reactions, containing no enzyme in both buffer systems, were also prepared as above.

Polyphosphate hydrolysis was tracked by running fractions of the reactions on TBE gels and staining polyphosphates present using toluidine blue O, as previously described (13). Briefly, reactions were diluted with Novex Hi-Density TBE Sample Buffer (Thermo Fisher LC6678), and run on precast TBE gels, either 20% (EC63155, Novex) for P45 and P65 reactions, or 6% (EC62655, Novex), for P700 reactions. Following electrophoresis, the gels were stained in 0.05% toluidine blue O in fixative solution (an aqueous solution of 25% methanol and 5% glycerol) for 15 minutes, with agitation, followed by 3 hours of washing in fixative solution and imaging. In concurrent analyses, released phosphate was quantified using BIOMOL GREEN phosphate detection kit as described above. All experiments were repeated at least three times with a minimum of four technical replicates per sample. Data was analyzed using PRISM 7 (GraphPad) software.

### Circular dichroism and *in silico* modeling of PRUNE1 mutations

PRUNE1 wild-type, D106N and R128Q muteins were analyzed by circular dichroism (CD) spectroscopy by collecting far-UV (<250nm) and near-UV (260 – 320nm) CD spectra. Prior to CD measurements, all samples were dialyzed twice in a base buffer, 20mM Tris-HCl, pH 7.8 to eliminate DTT that could potentially interfere with far-UV CD measurements. Following overnight dialysis, concentrations were determined by A280 measurements, using a theoretical extinction coefficient based on amino acid sequence of each protein. Near-UV CD measurements were obtained at 0.75mg/ml, 0.7mg/ml and 0.45mg/ml for wild-type, D106N and R128Q samples respectively. All samples were further diluted (two-fold) using 0.02µM filtered dialysate for far-UV CD measurements. Far-UV CD spectra were evaluated at 0.38mg/ml, 0.35mg/ml and 0.21mg/ml for wild-type, D106N and R128Q samples respectively. CD measurements were performed on a Jasco 1500 spectropolarimeter at 25°C using a 1cm cell (Jasco, Type J3) for near UV-CD and 0.1cm cell (Jasco, Type J10559) for far-UV CD. The raw data were baseline subtracted and converted to units of molar ellipticity to correct for protein concentration differences. Degree of spectral difference between wild-type and D106N or R128Q muteins was assessed using a spectral classification technique (TQ Analyst^®^). The reported TQ Analyst^®^ similarity match score indicates how closely the spectrum of each mutant matches that of the wild-type protein. Highly comparable far-UV CD spectra result in scores >92.6, whereas highly comparable near-UV CD give TQ similarity match scores >98.

The RCSB Protein Data Bank was searched for a structure with high sequence homology to human PRUNE1. *S. cerevesiae* cytosolic exopolyphosphatase (PPX1) (PDB ID: 2QB7), with a Blast e-value of 2.3E-19 and 25% sequence identity (45% similarity) was the best hit. Since the active site residues were conserved, we did not attempt to generate a PRUNE1 homology model. PPX1 structure (shown in Fig. 2A) was rendered by exchanging orthologous active site residues: Asp41 to Asn, Asp127 to Asn, H149 to Gln. Three bound phosphate moieties P_T_, P_E1_ and P_E3_ shown in ball-and-stick, and the two catalytic metal ions as M1 and M2 as pink spheres.

### Animal studies

*Prune1*^*-/-*^ mice were generated by replacing exon 2 (ENSMUSE00001210512) of the *Prune1* gene with a β-galactosidase (lacZ) reporter cassette using VelociGene technology (14). Mice carrying the deletion were genotyped by a loss of allele assay as described previously (15). Targeted, cassette-deleted heterozygous mice were bred to obtain desired genotypes. Homozygous and heterozygous knockout mice cohorts derived from F1 breeding were used for phenotypic evaluation. All mice used in this study were in a 75% C57Bl/6NTac, 25% 129S6/SvEvTac genetic background, and housed in a pathogen free environment. Autoclaved water and sterile mouse chow were provided *ad libitum*. All experimental protocols, anesthesia, imaging procedures and tissue sampling procedures performed in this study were approved by the Regeneron Pharmaceuticals Inc. Institutional Animal Care and Use Committee (IACUC).

### Anti-PECAM1 staining and imaging

Embryo processing, immunostaining and imaging were performed as described previously (16). Briefly, embryos were collected at embryonic day (E) 9.5 and fixed for 3 hours in 4% paraformaldehyde at 4ଌ. Prior to staining, endogenous peroxidase activity was quenched with 3% H_2_O_2_. Non-specific antibody binding was blocked by preincubating embryos in 10% normal goat serum (S-1000, Vector Labs) and 0.5% Bovine Serum Albumin (BP1600, Fisher Scientific). Embryos were stained overnight at 4ଌ with rat anti-PECAM1 antibody at 1:300 dilution (553370, Pharmingen). After primary antibody incubation, embryos were incubated overnight with donkey anti-rat HRP secondary antibody at 1:1000 dilution (Jackson ImmunoResearch, 712-036-153) followed by incubation with 488-tyramide reagent at 1:25 dilution made using Dylight 488 NHS ester (Thermo Scientific, 46402,). Images were acquired through optical sectioning with an ApoTome.2 on a ZEISS Axio Zoom. Extended Depth of Focus images were generated by Zen software. Imaging was performed using Leica MZFLIII stereozoom microscope quipped with a 0.5x objective and 1.0x camera lens, with a zoom between 4.0-6.3x. Alternatively, stained embryos were imaged using optical projection tomography (OPT) as described previously (16). For OPT, samples were embedded in low melting agarose and subsequently cleared using a mixture of benzyl alcohol and benzyl benzoate (1:2 ratio). Sample was rotated stepwise (with a 0.9° step size) for one revolution and a GFP1 filtered autofluorescence view and a Cy3 fluorescence view (captured using a Cy3 filter) were acquired using Retiga Exi CCD camera. Reconstruction of all image slices yielded a 3D volumetric rendering of the specimen as described previously (17).

### Statistical analysis

Statistical analyses were conducted using Prism 7 (GraphPad) software. Unpaired Student’s t-test was used unless indicated otherwise. Statistical significance reported when *p*<0.05. Sample sizes, the statistical test and significance are indicated in each Figure legend. Where appropriate sample sizes were estimated using resource equation method.

## Results

### Clinical description of NMIHBA patients

Family SZ51 is composed of two affected siblings (Patient 1 and Patient 2), an unaffected female sibling, and non-consanguineous parents of European ancestry. Familial genomic analyses of these two affected patients and available family members through whole exome sequencing identified, as previously reported, shared compound heterozygous variants in *PRUNE1 (*9). Both affected siblings inherited a missense c.383G>A; p.Arg128Gln variant from their unaffected father and a nonsense c.520G>T; p.Gly174* variant from their unaffected mother (Fig. S1). The unaffected sister is a heterozygous carrier for the paternal missense variant only.

Detailed clinical evaluation of both affected siblings was performed through the NIH Undiagnosed Diseases Program (UDP) at ages 4 years and 20 months, respectively. Both patients presented with profound developmental delay, intellectual disability, brain atrophy, seizures, and absent language. Additional features not documented in the initial report include optic atrophy, esotropia, scoliosis, gastrointestinal reflux (GERD), hypotonia and spasticity. Onset of seizures in both siblings occurred at 6 months of age; it was observed initially as infantile spasms that developed into gelastic complex partial myoclonic seizures in Patient 1. EEG showed slow waking background and slow spike waves with progression to multifocal epileptiform discharges. Additional detailed clinical information for these patients can be found in supplementary information (including Table S2 and S3).

Additionally, we performed detailed literature and clinical review of all 35 NMIHBA patients reported to date in order to better understand the phenotypic spectrum of NMIHBA (Table 1, Figure 1) (8-11, 18-20). Consistent features across the majority of patients include severe global developmental delay (94.59%), profound intellectual disability (91.89%) with absent language (83.78%) and brain abnormalities (86.48%) characterized primarily by cerebral and cerebellar atrophy, thin corpus callosum, and white matter changes. Hypotonia and spasticity are features observed in the majority of patients (97.29%), whereas microcephaly and seizures (64.86%) appear to be more variable than initially reported, as more cases are documented. Visual (43.24%) and gastrointestinal (37.83%) problems are additional clinical features in a sizable number of patients. Interestingly, Alhaddad et al, reported about half of their cohort having absent deep tendon reflexes (DTRs) and/or reduced nerve conduction velocities (NCVs) suggesting peripheral neuropathy as part of the clinical spectrum of NMIHBA(18). Other patients also showed neurogenic findings on electromyography (EMG) studies and spinal motor neuron involvement. However, the majority of patients were not assessed for these clinical phenotypes and the true prevalence of peripheral nervous system involvement in this disorder may be still underappreciated. A few patients, including probands described in this report developed scoliosis as well as foot anomalies such as talipes equinovarus or clubfoot, whether these features are secondary to the peripheral neuropathy and spinal muscular atrophy like phenotype or independent skeletal findings remains to be determined.

**Table 1.**
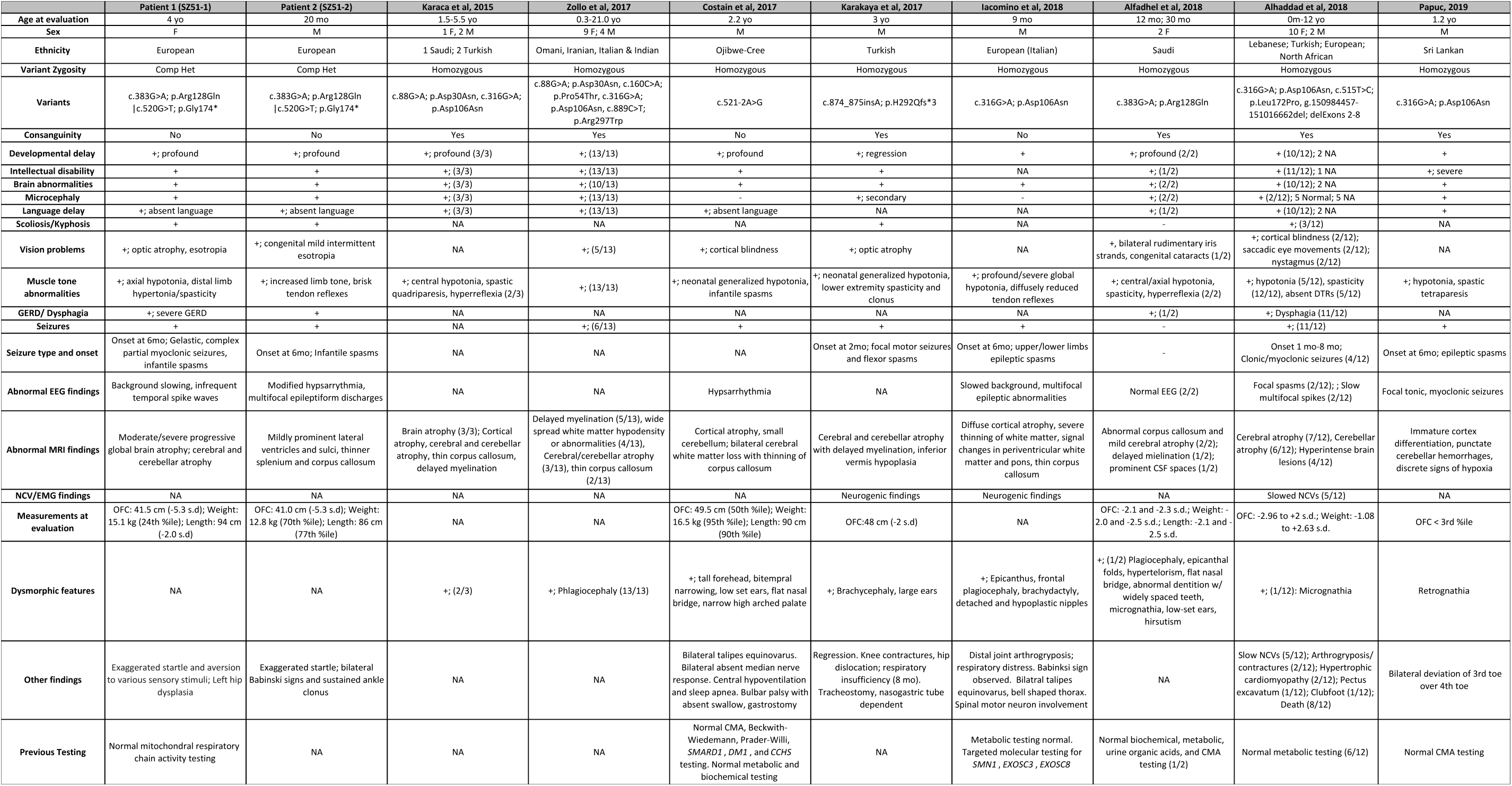
Summary of clinical findings of individuals with biallelic *PRUNE1* mutations. NCV: Nerve conductance velocity; EMG: Electromyography; OFC: Occipital frontal circumference; CMA: Chromosomal microarray testing; MR: Magnetic Resonance.

**Figure 1.**
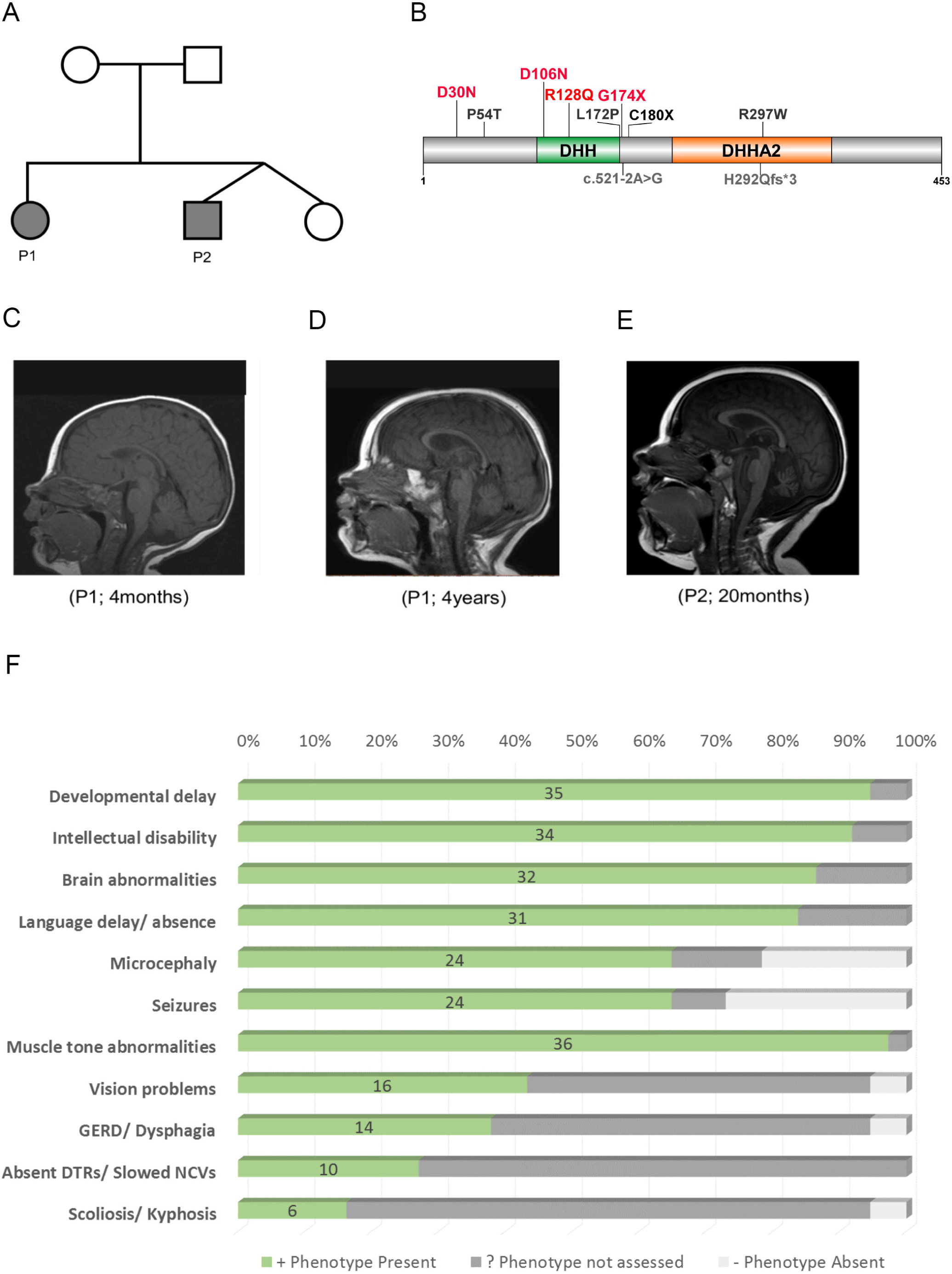
Clinical presentation of NMIHBA patients. (A) Immediate family pedigree of patient 1 (P1) and patient 2 (P2). (B) Pathogenic variants in *PRUNE1* identified in NMIHBA patients reported to date. The majority of pathogenic variants cluster in the DHH and DHH2 domains. Variants characterized in this study are highlighted in red. (C-E) Sagittal T1-weighted brain MRI images. (C) Image of patient 1 at 4 months of age showing no significant findings. (D) Image of patient 1 at 4 years of age showing severe cortical and cerebellar atrophy and milder corpus callosum and brainstem atrophy. The craniofacial ratio is decreased. The cerebellar vermis decreased in height from 29.5 to 25.6 mm between the time the two images were obtained. (E) Image obtained for patient 2 at 20 months of age demonstrating cerebellar vermis hypoplasia and mild cortical atrophy. (F) Frequency of clinical manifestations ascertained in reported NMIHBA cases.

**Figure 2.**
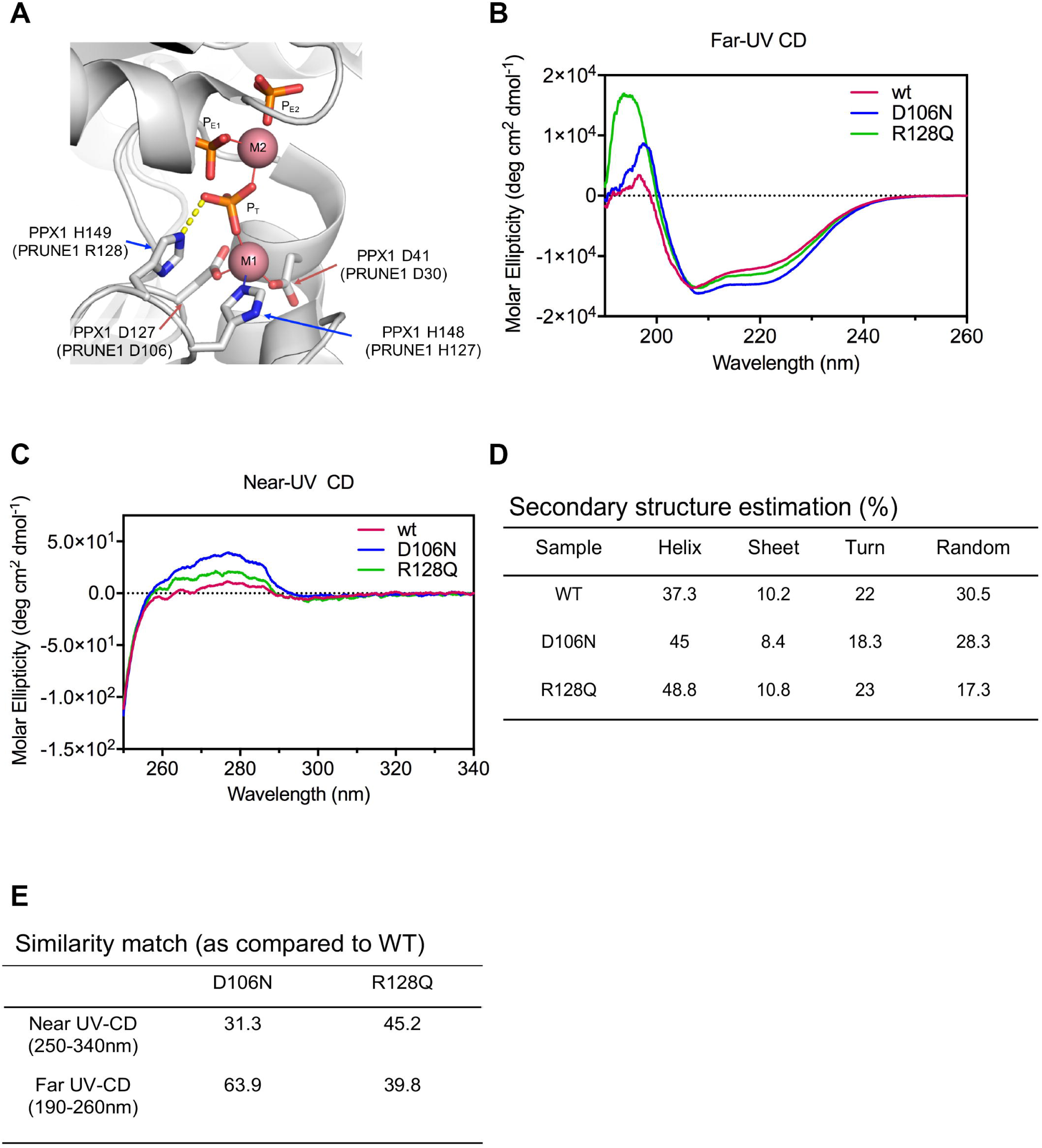
Missense mutations perturb secondary and tertiary structure of PRUNE1. (**A**) Homology modeling (based on 1.6 Å resolved *S*.*c. PPX1* structure, 2QB7) demonstrated D30, D106 and R128 residues fall within a metal-ion and phosphate binding interface representing the active-site for phosphate hydrolysis. Disruption of these charged residues within the active-site impair metal coordination (D30N and D106N) and substrate binding R128Q. Three bound phosphate moieties P_T_, P_E1_ and P_E3_ shown in ball- and-stick, and the catalytic metal ions (M1 and M2) as pink spheres. (**B**) Far-UV CD spectra reveals secondary structure differences between wild type and missense (D106N and R128Q) mutants. (**C**) Near-UV CD spectra signifies change in tertiary conformation of missense (D106N and R128Q) mutants as compared to the wild type. (**D**) Spectral deconvolution of far-UV CD spectra revealed differences in alpha-helical content between wild type and D106N or R128Q mutants. (**E**) Similarity match scores (based on quantitative assessment of similarity of far-UV and near-UV spectra) reveals low similarity of the missense mutants as compared to wild type.

Affected subjects have been reported to originate from countries around the globe (Figure S2). As expected, when homozygous alleles were identified there was often a history of consanguinity. Of note, all homozygous subjects have missense variant alleles – the only likely complete LoF null allele, c.G820T; p.G174*, is reported as a compound heterozygote with a missense variant suggesting some partial function is retained. The neurotypical phenotype of a heterozygous p.G174* carrier parents with presumed haploinsufficiency at the *PRUNE1* locus and the absence of either homozygous or compound heterozygous null alleles suggests that NMIHBA results from hypomorphic alleles or that null alleles may present with a different phenotype.

### Comparative modeling of PRUNE1 mutations

Similar to other members of the DHH superfamily, PRUNE1 contains a common N-terminal DHH domain resulting from four highly conserved motifs, with five invariant aspartates and two conserved histidine residues, one of which is replaced by Arginine (R128). One of the conserved aspartates is thought to act as nucleophile forming a charge-relay system with the conserved histidine, whereas the other aspartates bind divalent cations required for the catalytic activity (4). Interestingly, most of the pathogenic variants map to the highly conserved N-terminal DHH domain. In order to understand how NMIHBA-causing PRUNE1 variants drive the disease phenotype, we chose to characterize four variants (p.D30N, p.D106N, p.R128Q, p.G174*) within the conserved N-terminal domain for which we had patient-derived fibroblasts and/or comprehensive clinical information.

The importance of D30, D106, and R128 residues to PRUNE1 function is revealed by considering the crystal structure of a homologous enzyme, *S. cerevisiae* cytosolic exopolyphosphatase (PPX1) (PDB ID: 2QB7) (21). PPX1 is 25% identical and 45% similar to human PRUNE1 and has an equivalent enzymatic function (Fig. S3A)(4, 22). In this structure, a series of divalent metal ions (M1 and M2) and bound phosphate ions (P_T_, P_E1_ and P_E2_) delineate the active site and define residues necessary for binding the polyphosphate substrate as well as the catalytic metal ions. Strikingly, all three of the missense mutations under investigation map to the same region of the PPX1 active site: PPX1 residues D41 and D127 (equivalent to PRUNE1 D30 and D106) coordinate a divalent metal ion, while PPX1 residue H149 (equivalent to PRUNE1 residue R128) forms a charge-charge interaction with the adjacent bound phosphate (Fig. 2A).

Substitution of Asn for Asp (as in D30N and D106N) will eliminate that amino acid’s ability to coordinate the divalent metal ion, thereby abolishing catalytic activity and destabilizing the local protein structure that was ordered by interactions with that metal ion. Similarly, substitution of Arg 128 by Gln eliminates the positive charge on that residue, likely preventing the mutated residue from interacting with the negatively charged phosphate group of a bound substrate.

To further verify these structural predictions, we performed circular dichroism (CD) on mammalian cell (Expi293) derived recombinant muteins harboring the D106N and R128Q variants. Whereas D106N and R128Q recombinant proteins expressed and migrated similarly to wild-type, rapid turnover of the D30N mutant precluded protein purification and further analyses (Figure 3A; Fig. S3B and C). Far and near-UV CD spectra were collected for wild-type PRUNE1 and the two mutants (D106N and R128Q). Qualitative differences in the far and near-UV CD spectra were observed for each mutant compared to wild-type PRUNE1 (Fig 2B and C). Spectral deconvolution of the far-UV CD revealed differences in alpha-helical content in the D106N and R128Q muteins (45% and 48.8% respectively) as compared to wild-type (37.3%). Near-UV CD analyses revealed differences in tertiary structure between D106N and R128Q muteins as compared to wild-type. Furthermore, increased magnitude of CD signal particularly in the region contributed by Phe and Tyr residues indicated change in overall tertiary structure. The increase in CD signal possibly resulted from a more rigid conformation of the aromatic sides chains induced by the D106N and R128Q mutations. Lastly, quantitative assessment of spectral similarity (assessed by TQ similarity match scores) of the far and near-UV CD between the wild-type and muteins demonstrated a low similarity between the spectra of the mutants compared to that of wild-type (Fig 2D and E). Therefore, our preliminary *in silico* and protein biochemistry analyses predicted that these three missense mutations could result in loss of protein function directly, by abolishing substrate binding and catalytic activity, and furthermore, by destabilizing the structure of the active site.

**Figure 3.**
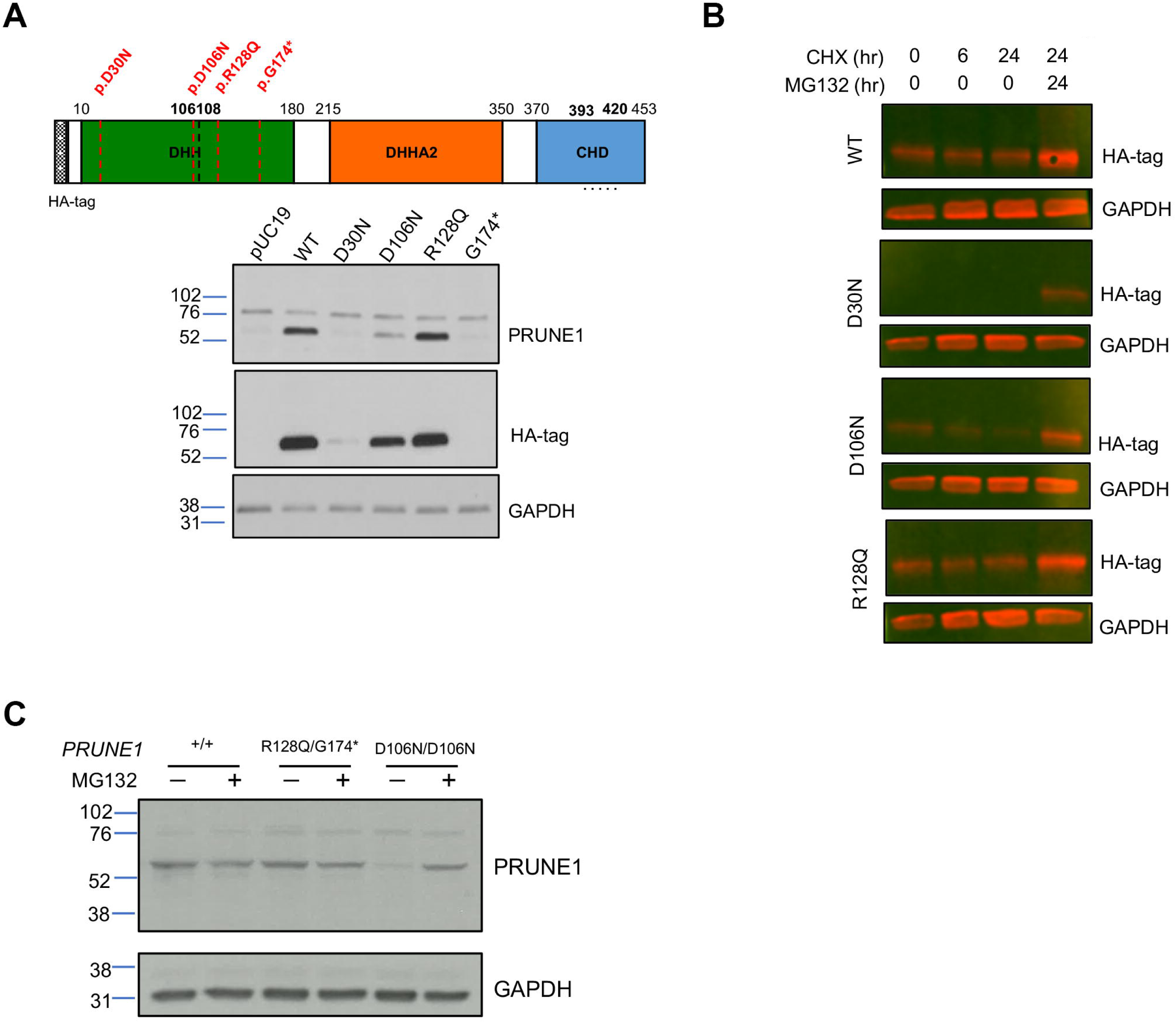
D30N and D106N variants result in reduced protein stability and proteosomal degradation, whereas R128Q variant results in stable mutant protein. (**A**) HEK293 cells were transfected with N-terminal HA-tagged wild type or mutant PRUNE1 cDNA. Equal amounts of protein from whole cell lysate were used for immunoblotting using antibodies against N-terminal HA-tag, and C-terminal PRUNE1 epitope (a.a. 393-420, dotted line). GAPDH was used as a loading control. G174*mutant showed no expression, whereas D30N and D106N mutants showed significantly reduced expression as compared to wild type and R128Q levels. (**B**) Immunoblots of cells overexpressing N-terminal HA-tagged wild type or mutant PRUNE1 treated with cycloheximide (CHX) at 200µM for 0, 6 and 24 hours (with or without proteasome inhibitor MG132 at 15 µM). (**C**) Immunoblotting of endogenous PRUNE1 in patient derived fibroblasts harboring compound heterozygous R128Q; G174* variants or homozygous D106N missense variant treated with or without MG132(15 µM).

### Missense mutations result in loss of exopolyphosphatase activity

In order to evaluate protein stability, mammalian expression constructs encoding epitope (HA)-tagged wild type or missense PRUNE1 muteins (D30N, D106N, R128Q, G174*) were transfected into HEK293 cells (Fig S4A and B). HEK293 cells have no detectable endogenous PRUNE1 expression. At steady state, immunoblotting analyses of whole cell lysates demonstrated a significant reduction of PRUNE1 levels in cells expressing D30N or D106N variants in comparison to wild type tagged protein. Whereas lysates from cells expressing G174* showed no discernable protein product, the R128Q variant-expressing cells showed similar expression compared to cells expressing the wild type protein (Fig. 3A). Furthermore, cycloheximide chase assay demonstrated rapid and significant loss of D30N and D106N mutant proteins, that was rescued by the cell-permeable proteasome inhibitor, MG132 (Fig 3B). In contrast, the R128Q variant and wild type PRUNE1 levels were stable after synthesis, as they were observed at >60% of baseline (0hr) levels after 24hr of cycloheximide treatment (data not shown). In subsequent experiments we evaluated PRUNE1 protein levels in patient-derived skin fibroblasts harboring the compound heterozygous R128Q/G174* variants or homozygous D106N variants (Fig. S4B). PRUNE1 levels were comparable between fibroblasts harboring compound heterozygous variants and the wild type allele. However, cells homozygous for the D106N variant showed dramatic reduction in PRUNE1 levels at steady state. Proteasome inhibition by MG132 rescued the loss of the mutant protein in these cells (Fig. 3C). Taken together, these studies suggested that the D30N and D106N variants lead to reduced protein stability and degradation by the proteasome pathway, whereas the R128Q variant results in a stable mutant protein.

PRUNE1 has been ascribed exopolyphosphatase and phosphodiesterase activity based on sequence homology to yeast exopolyphosphatase PPX1 and bacterial nuclease RecJ respectively (4). In subsequent experiments we assessed how missense variants in PRUNE1 influence its enzymatic function. GloSensor assays established no detectable cAMP or cGMP phosphodiesterase activity of wild type or missense PRUNE1 variants as compared to PDE4A and PDE5A (Fig S5A). However, consistent with the findings of Tammenkoski et al.(22), wild type PRUNE1 catalyzed polyphosphate hydrolysis of short-chain polyphosphates P_3_ (*k*_*cat*_ = 6.24s^-1^; *K*_*m*_ = 8.4µM; *V*_*max*_ = 0.06 µM/s) and P_4_ (*k*_*cat*_ = 4.2s^-1^ ; *K*_*m*_ = 16.13µM; *V*_*max*_ = 0.04 µM/s) using Mg^2+^ as the metal cofactor (Fig 4A and B), and demonstrated no expolyphosphatase against long chain polyphosphates P_45_, P_65_ and P_700_ (Fig S5C). D106N and R128Q variants completely abolished short-chain exopolyphosphatase compared to wild type as assessed by BIOMOL®Green phosphate reagent using P_3_ (1-50 µM) and P_4_ (5-100 µM) as polyphosphate substrates (Fig 4C). Similarly, *E*.*coli* expressed D106N and R128Q variants had no detectable short-chain exopolyphosphatase activity as compared to wild-type (Fig S5B).

**Figure 4.**
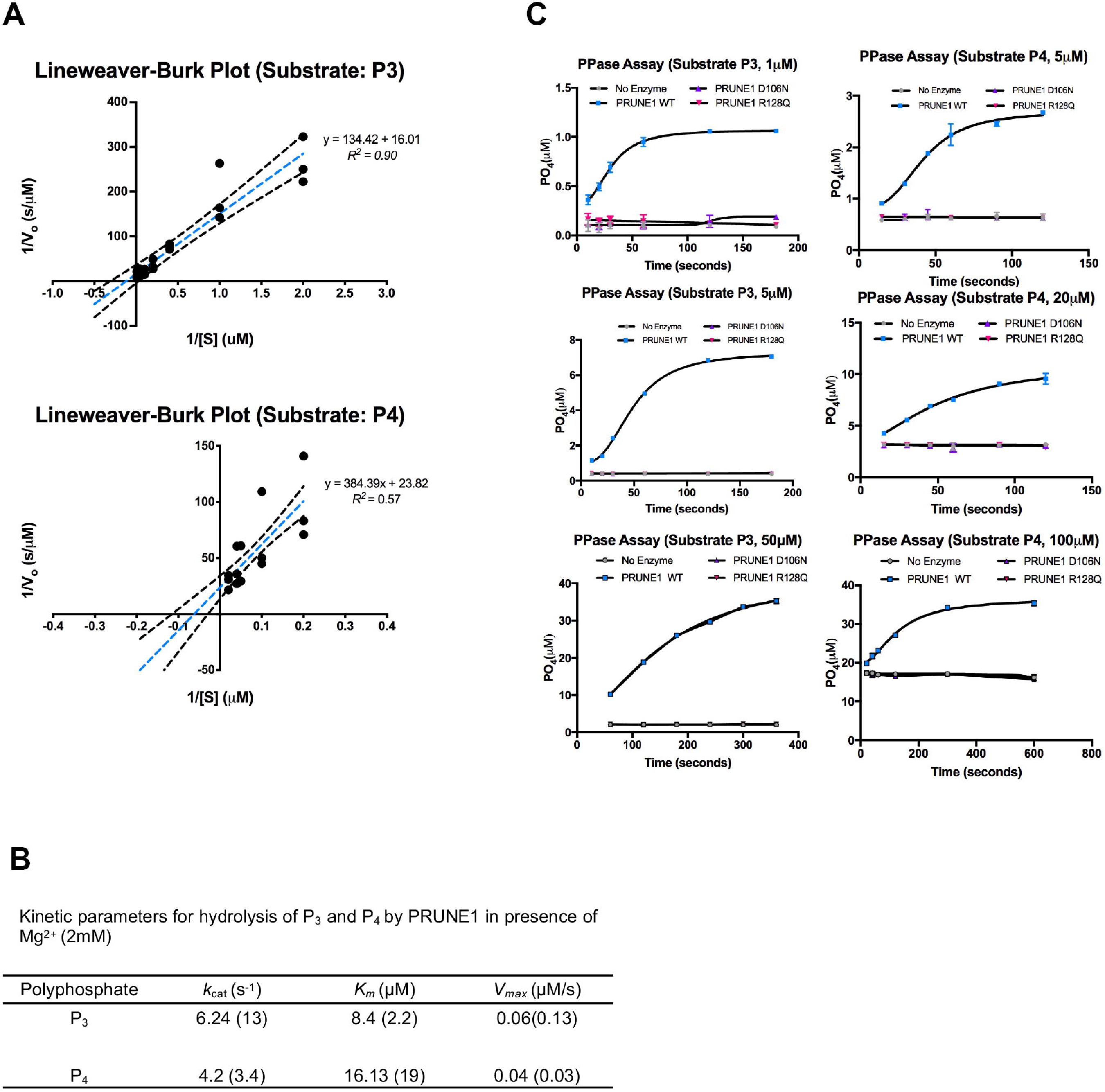
D106N and R128Q variants result in loss of short chain exopolyphosphatase activity. (**A)** Lineweaver-Burk plots depicting short-chain exopolyphosphatase kinetics of wild type PRUNE1 using sodium tripolyphosphate (P3) and sodium tetrapolyphosphate (P4) as substrates. Dotted black lines represent 95% CI. (**B**) Kinetic parameters for hydrolysis of polyphosphates (P3 and P4) by wild type PRUNE1 in the presence of Mg^2+^ (2mM) as the cofactor. Values reported by Tammenkoski et al. are shown within parenthesis(22). (**C**) Short-chain exopolyphosphatase activity of wild type, D106N and R128Q mutants on P3 and P4 determined using fixed-time BIOMOL Green phosphate detection assay. Data represented as mean ± SEM over 3 independent experiments with 6 technical replicates per sample.

Taken together, these results indicated that missense variants D30N, D106N and R128Q result in loss of PRUNE1 function either due to impaired protein stability or due to the loss of enzymatic function. These findings are consistent with the observed phenotypic concordance between patients harboring homozygous missense D30N, D106N variants and the compound heterozygous probands (p.Arg128Gln; p.Gly174*) described herein (patient 1 and patient 2). Importantly, these studies support loss of PRUNE1 function as the molecular etiology of NMIHBA.

### Loss of Prune1 results in embryonic lethality in mouse

In an effort to model NMIHBA *in vivo*, we next studied the consequences of *Prune1* loss-of-function by generating *Prune1*^*-/-*^ mice as shown in Fig S6A and B. Here homologous recombination was employed to replace exon 2 with a *lacZ* reporter cassette. A stable transcript derived from this allele predicts a non-functional Prune1 protein missing amino acids 15-454. Importantly, this mutant allele provides for a *lacZ* reporter, enabling spatio-temporal localization of *Prune1* gene reporter activity. *Prune1*^*+/-*^ animals (both sexes) are viable and appeared normal as compared to wild-type littermates, with no differences in body weight, body composition, serum chemistry and hematology (data not shown). *LacZ* expression profiling at E12.5 demonstrated widespread expression across multiple tissues including the brain (Fig S6B and D). In adult *Prune1*^*+/-*^ mice (postnatal day 56) strong CNS expression was observed in the hippocampus, cerebellum, amygdala, hypothalamus and the cortex (Fig S6D).

No homozygous mutants were found among 55 pups at postnatal day 10 from *Prune1*^*+/-*^ intercrosses (Fig S6C). In light of this presumed embryonic lethality, embryos were collected at various developmental time points. Mendelian ratios were maintained at E9.5 (Fig S6C), however no *Prune1*^*-/-*^ embryos were found beyond E12.5 suggesting lethality between E9.5 and E12.5. Whole mount examination of embryos and yolk sacs at E9.5 (∼25 somites) employing endothelial-specific Pecam 1 immunochemistry revealed significant defects in vessel morphology in Prune1 mutants. First, *Prune1*^*-/-*^ yolk sacs exhibit a profound decrease in capillary plexus formation and absence of differentiated vessels, indicative of perturbed vascular remodeling (Fig S7B). Second, mutant embryos also show severe defects in formation and remodeling of the cephalic vascular plexus, most strikingly revealed in 3D in Optical Projection Tomography (OPT). In addition, anomalies to the limb bud vasculature, dorsal vascular plexus (along the spinal cord), as well as marked cardiac hypoplasia were also noted in all *Prune1*^*-/-*^ embryos (Fig S7A, Fig 5A and B).

**Figure 5.**
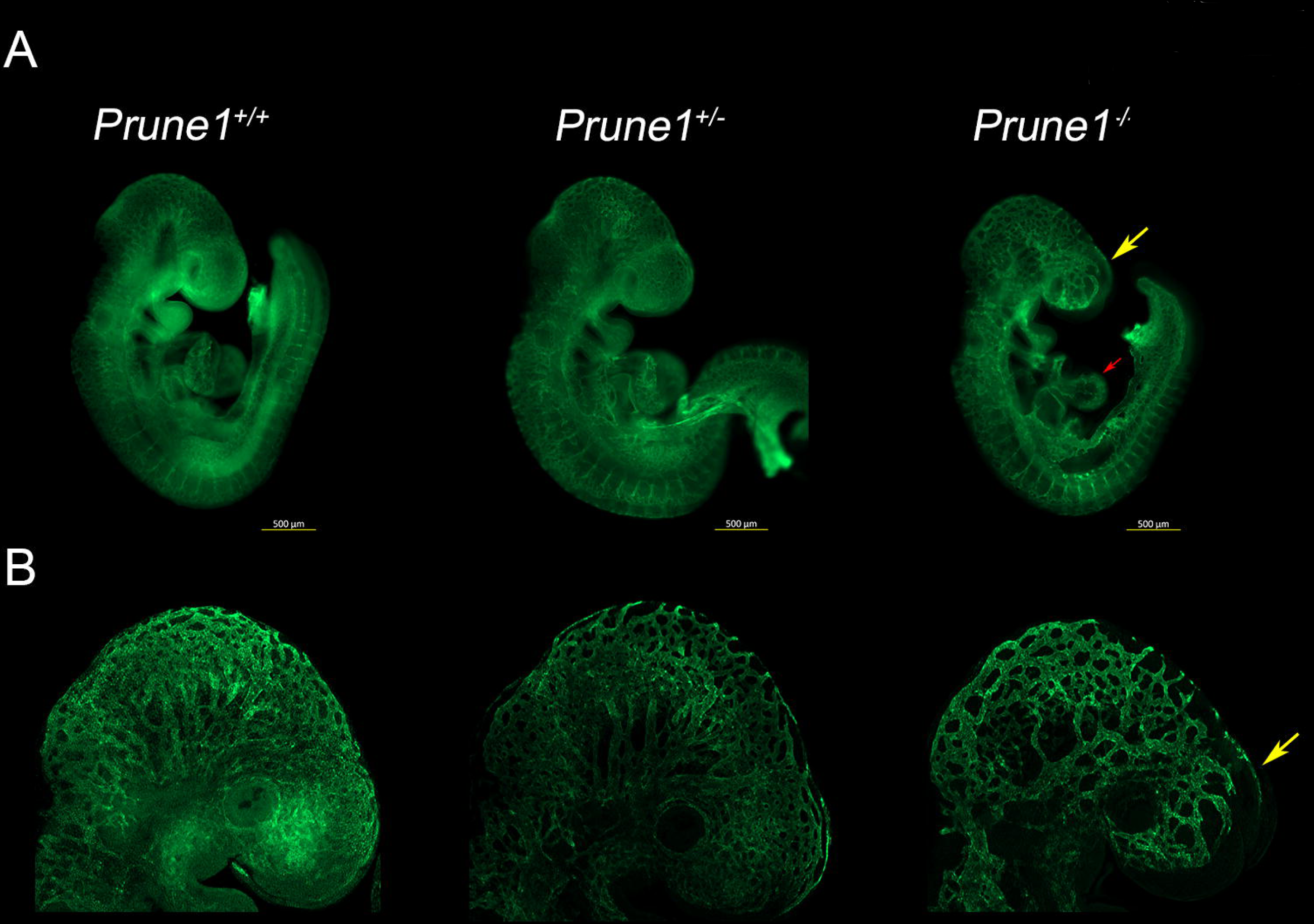
Loss of *Prune1* results in vascular defects with significant disruption of the cephalic vascular plexus. (**A**) Representative whole mount Pecam1 staining at E9.5 demonstrated reduced plexus branching and perturbed capillary sprouting within the cephalic region (frontonasal prominence and brain, yellow arrow) in the *Prune1*^*-/-*^ embryos as compared to wild type and heterozygous littermates. Moreover, *Prune1*^*-/-*^ embryos displayed cardiac defects observed as a less intricate appearance of the endocardium when compared to littermate controls (red arrow) (**B**) Higher magnification (6.3X) images further highlight disruption of the cephalic vascular plexus. A total of 3 *Prune1*^*+/+*^, 2 - *Prune1*^*+/-*^ and 5 *Prune1*^*-/-*^ embryos were analyzed in two independent experiments.

## Discussion

Since our initial report of *PRUNE1* as a novel candidate gene for abnormal neurodevelopment and brain malformation, 37 NMIHBA patients have been reported to date by us and others (8-11, 18-20, 23, 24). Detailed clinical and literature review of these reported NMIHBA cases allows for a more comprehensive view of this novel neurodevelopmental disorder (Table 1, Figure 1). The main features defining the disorder, namely profound developmental delay, intellectual disability and brain malformations, remain consistent; however additional clinical features such as optic nerve atrophy, gastrointestinal problems and peripheral nervous system involvement have emerged with additional reports of patients with NMIHBA. Subsequent reporting of patients with novel genetic disorders after the initial report, enables better understanding of their phenotypic spectrum and may help inform on disease progression and natural history when new patients are diagnosed with these conditions.

The worldwide distribution of reported patients and disease associated variants conforms with the clan genomics hypothesis where rare alleles of high effect arising in recent ancestors are more likely to contribute to disease burden versus common variants of small effect in the population (25). These rare alleles are more likely to come together in homozygosity in certain populations due to genetic mechanisms such as drift in founder populations or inbreeding in consanguineous populations. For example, the splicing variant allele c.521-2A>G has now been reported multiple times as a founder allele in the Cree population from Canada (19, 26). Similarly, the D30N allele, so far reported in Saudi and Omani individuals (8, 9), appears to be a founder allele in Middle Eastern populations; whereas other pathogenic rare alleles have been reported once so far in single families from different populations around the world. Interestingly, the most common pathogenic allele reported in NMIHBA patients is the D106N variant, observed in individuals from diverse ethnic backgrounds from Turkey, Italy, Lebanon, Sri Lanka, and Japan. While this variant appears to be a founder allele in the Turkish population, the diversity of populations where it has been observed poses the possibility that the D106 residue is a recurrent mutation site and that the allele has arisen multiple times in different population haplotypes.

Furthermore, in this study we investigated the molecular etiology underlying PRUNE1 dysfunction in NMIHBA. Based on our *in vitro* and *in vivo* data, we conclude that the molecular pathomechanism of this rare genetic disorder is reduced or altered function of PRUNE1 due to hypomorphic alleles. The majority of disease associated alleles including the splice site variant within intron 4 (c.521-2A>G: IVS4-2A>G) cluster in the catalytic DHH domain of PRUNE1, underscoring the critical role of this conserved domain in NMIHBA pathogenesis. Using *in silico* and biochemical approaches we explored the functional effects of four reported pathogenic *PRUNE1* variants: p.D30N, p.D106N, p.R128Q and p.G174* on PRUNE1 function and examined potential developmental and organismal functional consequences of homozygous *Prune1*^*-/-*^ ablation.

Protein homology modeling using the *S*.*cerevisiae* orthologue, PPX1 demonstrated that the D30, D106 and R128 residues fall within a metal-ion and phosphate binding interface representing the active-site for phosphate hydrolysis. Disruption of these charged residues within the active-site is predicted to impair metal coordination (D30N and D106N) and substrate binding (R128Q). Overexpression of human *PRUNE1* cDNA harboring NMIHBA pathogenic variant alleles in HEK-293 cells demonstrated a complete loss of protein product for the D30N mutant and a dramatic reduction in the D106N product. Both isoforms were rescued upon inhibition of the proteasome degradation pathway, suggesting that the loss of metal-binding residues destabilizes local protein structure resulting in protein degradation. In contrast, the R128Q variant was stably expressed at levels comparable to wild-type protein. Patient derived fibroblasts further confirmed these findings, implicating the metal coordinating D106 residue to be critical for maintaining active-site structure and protein stability. It has been proposed that the R128 (H149 in *S*.*c*.-PPX1) along with the binuclear metal cluster (M1 and M2 in Fig 2A) coordinate binding of one of the phosphate moieties (P_T_; Fig 2A) of the substrate, rendering the P-O bond more susceptible to nucleophilic attack. This model would suggest impaired polyphosphate processivity by the R128Q mutant. Consistent with such a model, *in vitro* short-chain exopolyphosphatase assays using recombinant proteins derived from mammalian and *E*.*coli* overexpression systems demonstrated loss of exopolyphosphatase activity in R128Q and D106N mutants as compared to the wild-type. Kinetic parameters for hydrolysis of short-chain polyphosphates P_3_ and P_4_ (in the presence of 2mM Mg2^+^) by wild-type PRUNE1 in our assays were comparable to those reported by Tammenkoski *et al*.(22). Our data related to the R128Q mutant in conjunction with enhanced kinetics of P_3_ hydrolysis by R128H mutant (where the positive charge is retained) demonstrated by Tammenkoski *et al*., further reinforce a model wherein binding of the metal ion at the active site accelerates substrate binding, and the bound substrate in turn enhances the enzyme affinity for the metal ion (22). Therefore, mutations that disrupt metal coordinating residues or substrate binding residues may result in loss of enzymatic function. Furthermore, in line with previous reports we demonstrate that wild-type PRUNE1 has no detectable cyclic nucleotide (cAMP or cGMP) phosphodiesterase activity or long-chain exopolyphosphatase activity (22). Taken together, these studies suggest loss of short-chain exopolyphosphatase activity as a potential mediator of NMIHBA pathophysiology. In light of our biochemical observations on disease associated mutant proteins, it is intriguing to note that Zollo et al.(8), ascribe enhanced short-chain exopolyphosphate activity (based on P_4_ hydrolysis) for two NMIHBA variants they characterized including the D30N mutant which is demonstrably a loss of function variant in our analyses. Furthermore, their reported K_cat_/K_m_ ratio for P_4_ hydrolysis by wild-type PRUNE1 (0.014µM^-1^s^-1^) is an order of magnitude different than those observed in the present study (0.26µM^-1^s^-1^) and reported by Tammenkoski et al. (0.18µM^-1^s^-1^) (8, 22). These discrepant results may be due to differences in cofactor concentration (10mM Mg2^+^ as opposed to 2mM Mg2^+^ in our assays and Tammenkoski et al.), substrate purity, or other confounding factors specific to their experimental methods used.

At the time of our initial discovery of *PRUNE1* as the disease causing gene, no mouse or other model organism data were available to provide further insights into the biological role of PRUNE1 in neurodevelopment and disease pathophysiology. Conservation at the protein level and similar developmental expression suggested preserved function between the mouse and human orthologues. Based on functional analyses of disease causing human alleles, we engineered a mouse model of NMIHBA by knocking out the murine *Prune1* gene. Since the original publication of *PRUNE1* as a novel disease gene in Karaca et al (9), a *Prune1*-null mouse was developed and phenotyped by the International Mouse Phenotyping Consortium (IMPC). (http://www.mousephenotype.org/data/genes/MGI:1925152). Consistent with the IMPC data, we observed fully penetrant embryonic lethality in the null mouse prior to E12.5. Our in depth developmental phenotyping of the *Prune1*-null embryos demonstrated profound vascular defects manifested by poorly branched vessels within the yolk sac, cardiac hypoplasia and a disrupted cephalic vascular plexus resulting in embryonic lethality between E9 – E10. However, our analyses do not address if the observed vascular anomalies result from altered hemodynamics secondary to impaired vasculogenesis within the yolk sac or if these represent cell-autonomous effects driven by loss of Prune1.

*LacZ* reporter expression in heterozygous mice demonstrates broad diffuse *Prune1* expression across multiple tissues and cell lineages at E12.5 (Fig. S6B). Consistently, *Prune1* expression in adult mice is observed across multiple tissues with strong expression within the CNS (Fig. S6D). Despite this widespread expression profile, the most overt phenotype arising from loss of *Prune 1* are vascular deficits most prominent in the cephalic vascular plexus possibly resulting from a local deficit in angiogenic cues or due to altered hemodynamics downstream of defective vascular development. Further analyses are required to discriminate the effect of hemodynamic defects versus the signaling perturbations downstream of Prune1 loss of function, and more importantly as to why the cephalic vessels are most affected.

It is intriguing as to why homozygous deletion results in embryonic lethality in mouse, while probands in this report and others survive past gestation. Whereas two patients have been reported to be homozygous for splice-site and frameshift variants in *PRUNE1* (11, 20), comprehensive analyses of protein-coding variation in *PRUNE1* across several large scale human genetic variation databases including ExAC/gnomAD and our internal (DiscovEHR) database identified no homozygous carriers of predicted loss-of-function variants despite the observation that heterozygous loss-of-function variants are well tolerated (Fig. S8) (27, 28). Residual enzymatic function of the hypomorphic missense alleles beyond the range of detection of our *in vitro* assays, alternative non-enzymatic developmental function (in mouse) or potential compensation by other phosphatases during neurodevelopment (in human) could potentially explain the observed species-specific difference in phenotype. Future analyses using knock-in mouse models may help discern the allelic architecture of NMIBHA and allow for functional interrogation of PRUNE1 in mammalian development.

Polyphosphate has been implicated in diverse physiological processes in higher eukaryotes, including apoptosis, mTOR activation and neuronal signaling (29-31). This pleiotropy is consistent with the essential role of inorganic diphosphate (PP_i_) for cellular metabolism across taxa and the potential role of polyphosphate metabolizing enzymes (including PRUNE1) in maintaining phosphate homeostasis. However, mechanistic understanding of how polyphosphate metabolism influences cellular metabolism and in turn organogenesis remains obscure. More recently, Cremers et al. proposed polyphosphate as a conserved modifier of amyloidogenic processes capable of mitigating neurotoxicity induced by amyloid fibers (32). Further studies using mouse models that closely recapitulate the NMIHBA phenotype will help uncover mechanisms by which PRUNE1 influences neurodevelopmental and neurodegenerative processes. The fact that NMIHBA-causing *PRUNE1* variants lack exopolyphosphatase activity may facilitate the exploration of therapeutic avenues that would allow for preemptive rather than symptomatic intervention of NMIHBA in the future.

## Acknowledgements

We wish to thank the patients and families that consented to be part of this study and acknowledge Kristy Neiman and William Poueymirou for their assistance with mouse colony management. This study was supported in part by the joint National Human Genome Research Institute (NHGRI) / National Heart Blood and Lung Institute (NHLBI) grant UM1 HG006542 of the National Institutes of Health to the Baylor-Hopkins Center for Mendelian Genomics and NINDS R35 grant to JRL.

## Supplementary Methods

**Figure S1.**
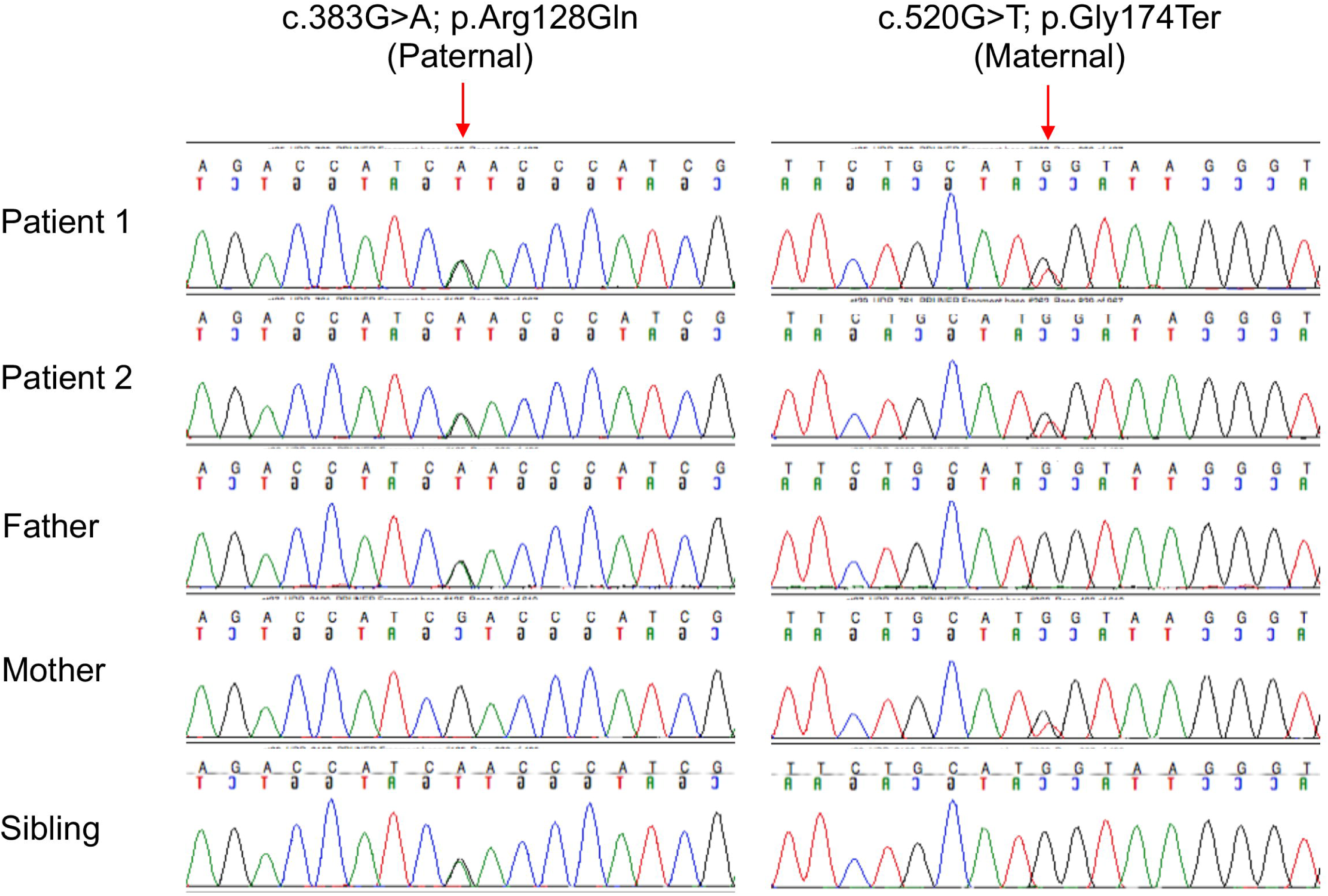
Chromatograms showing the familial segregation of the compound heterozygous variants inherited by patient 1 and patient 2.

**Figure S2.**
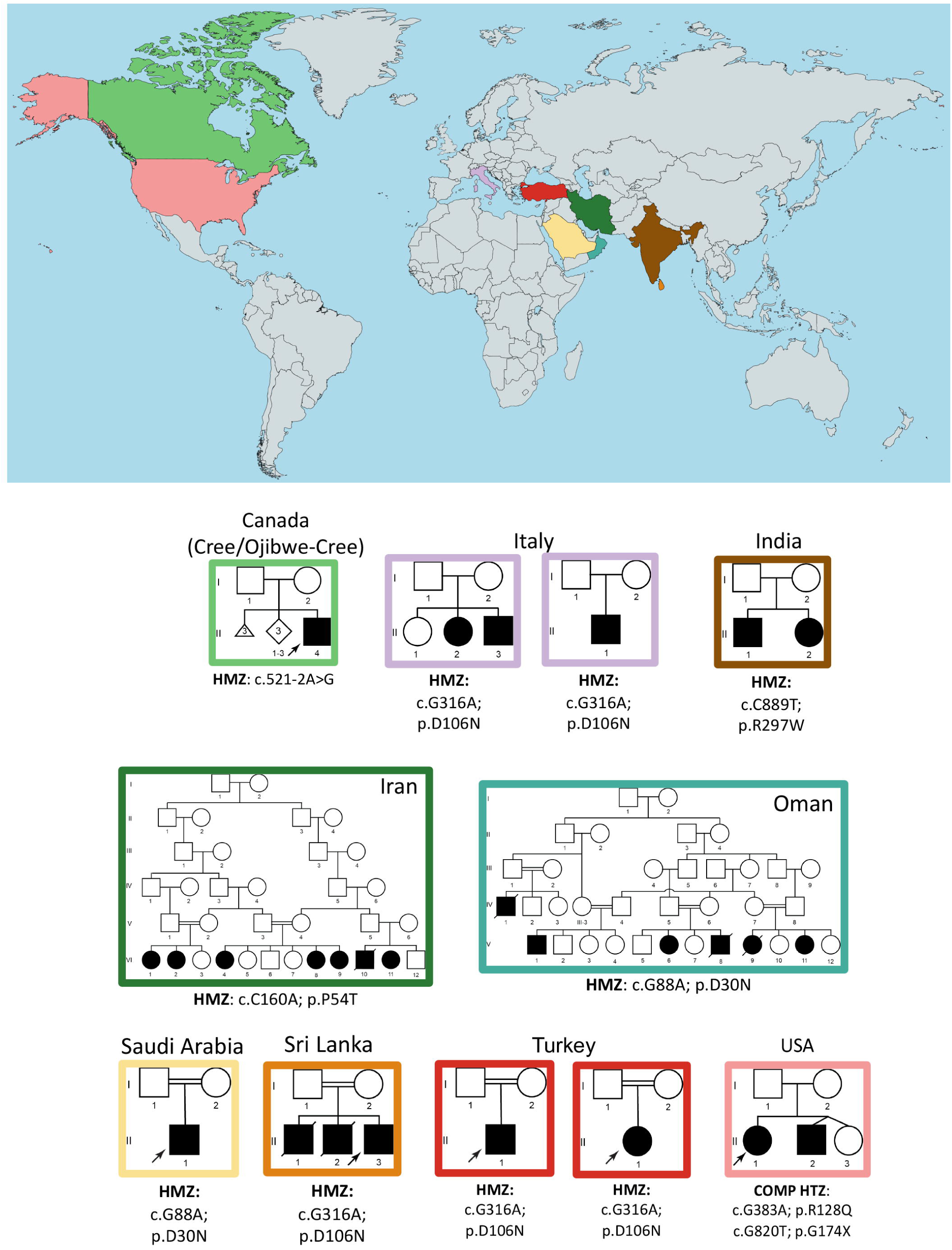
World-wide distribution of identified pathogenic variant *PRUNE1* alleles. HMZ: Homozygous. COMP HTZ: Compound heterozygous.

**Figure S3.**
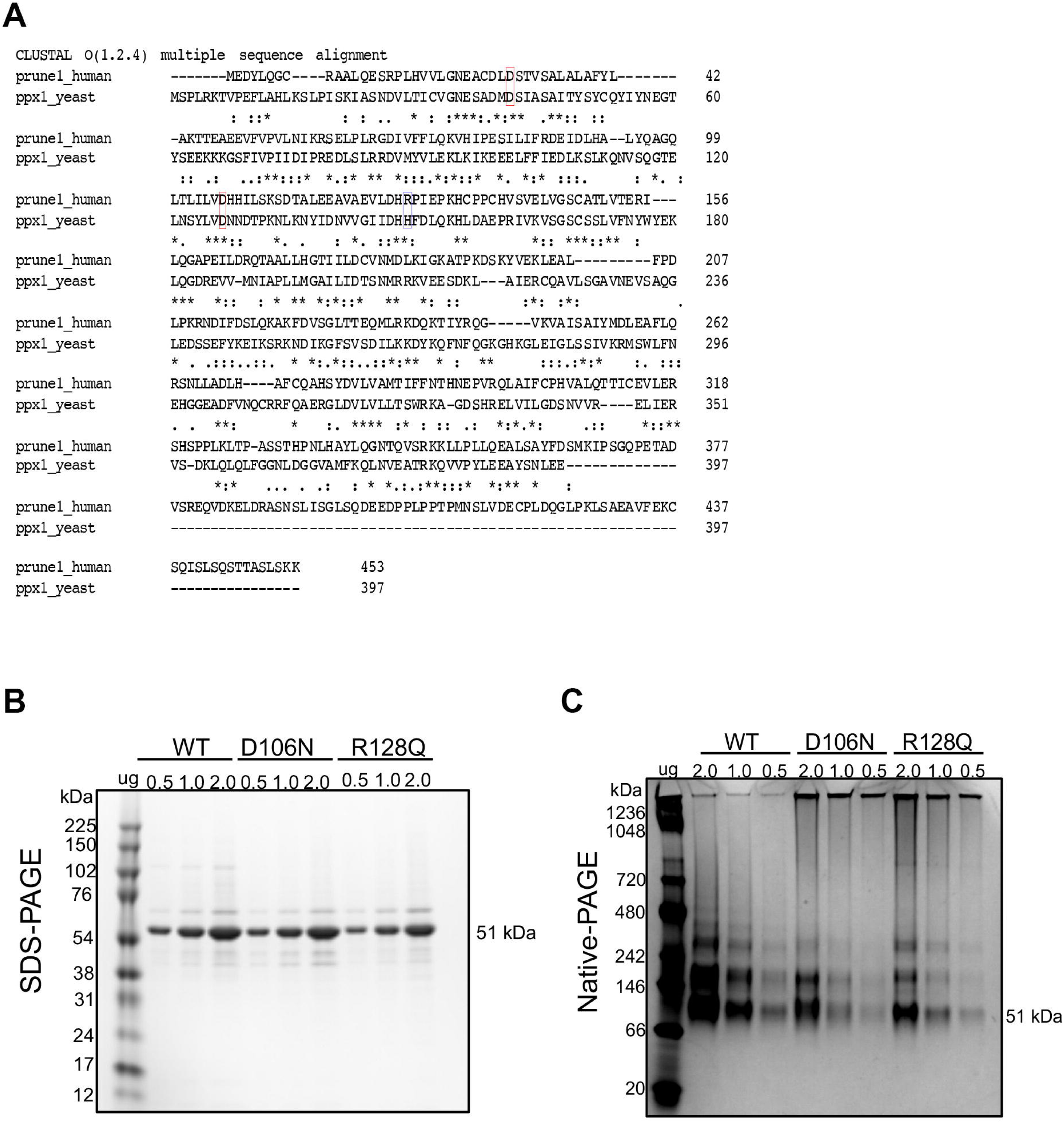
(**A**) Sequence alignment of *S*.*c*. PPX1 and PRUNE1. PPX1 is 25% identical and 45% similar to PRUNE1 and has equivalent enzymatic function. Active site residues are conserved between human and yeast enzymes. PPX1 residues D41 and D127 (corresponding to PRUNE1 D30 and D106, red box) coordinate divalent metal ion, while PPX1 residue H149 (equivalent to PRUNE1 R128, blue box) results in charge-charge interaction with adjacent bound phosphate. (**B and C**) SDS-PAGE and Native-PAGE of Expi-293 derived wild type, D106N and R128Q proteins.

**Figure S4.**
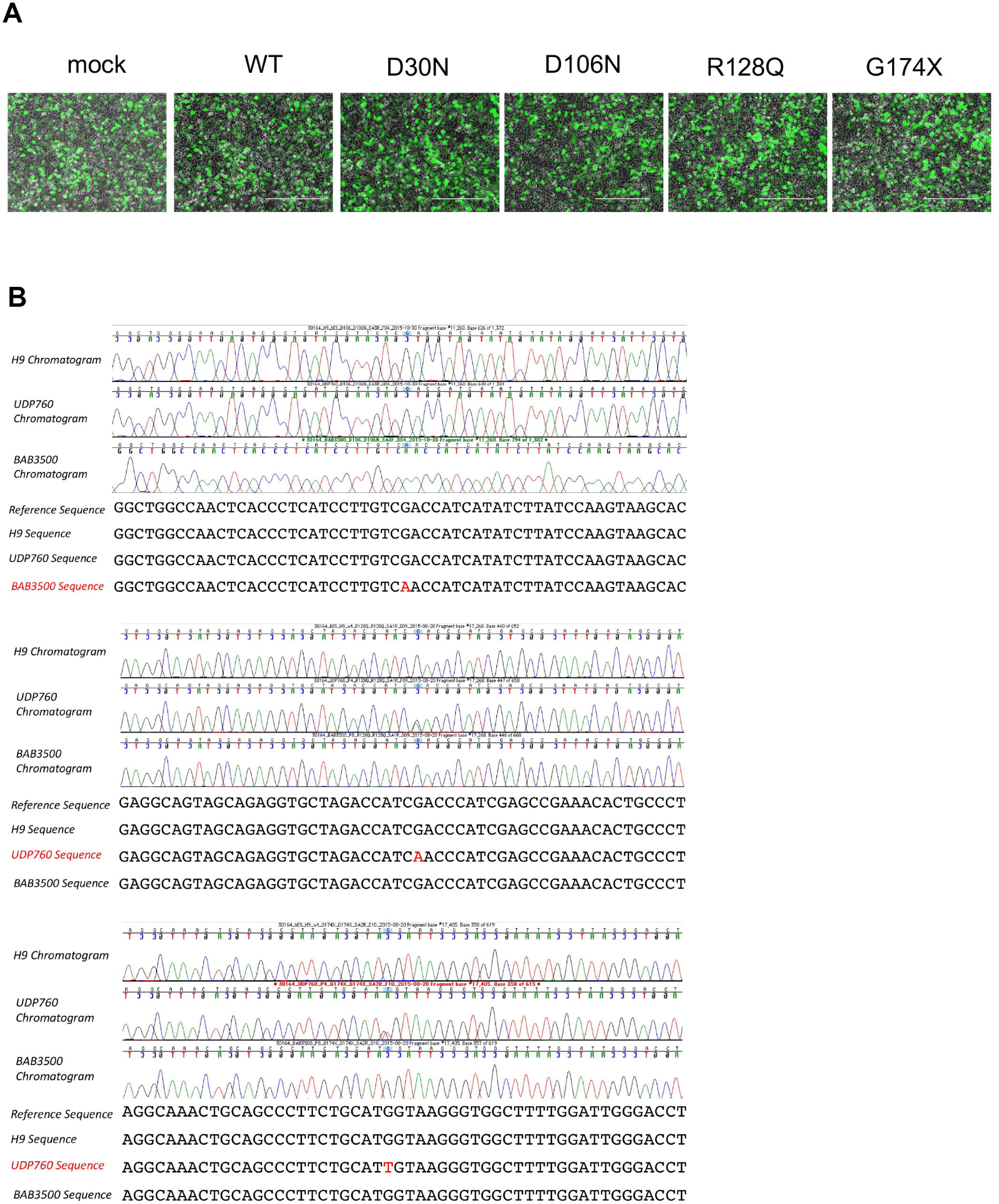
(**A**) GFP expression (a mammalian expression construct under CMV promoter) upon co-transfection with N-terminal HA-tagged wild type or mutant PRUNE1 cDNA in HEK293 demonstrating equivalent transfection efficiencies (Fig. 3A). Fluorescence microscope images of transfected cells demonstrate similar transfection efficiency across all samples. (**B**) Sanger sequencing of patient-derived fibroblasts. Chromatograms showing c.316G>A (p. D106N) and compound heterozygous variants c. 383G>A (p.Arg128Gln), c.520G>T (p.Gly174*) identified in BAB3500 and UDP760 cells respectively. Genomic DNA from H9 ES cells was used as normal control.

**Figure S5.**
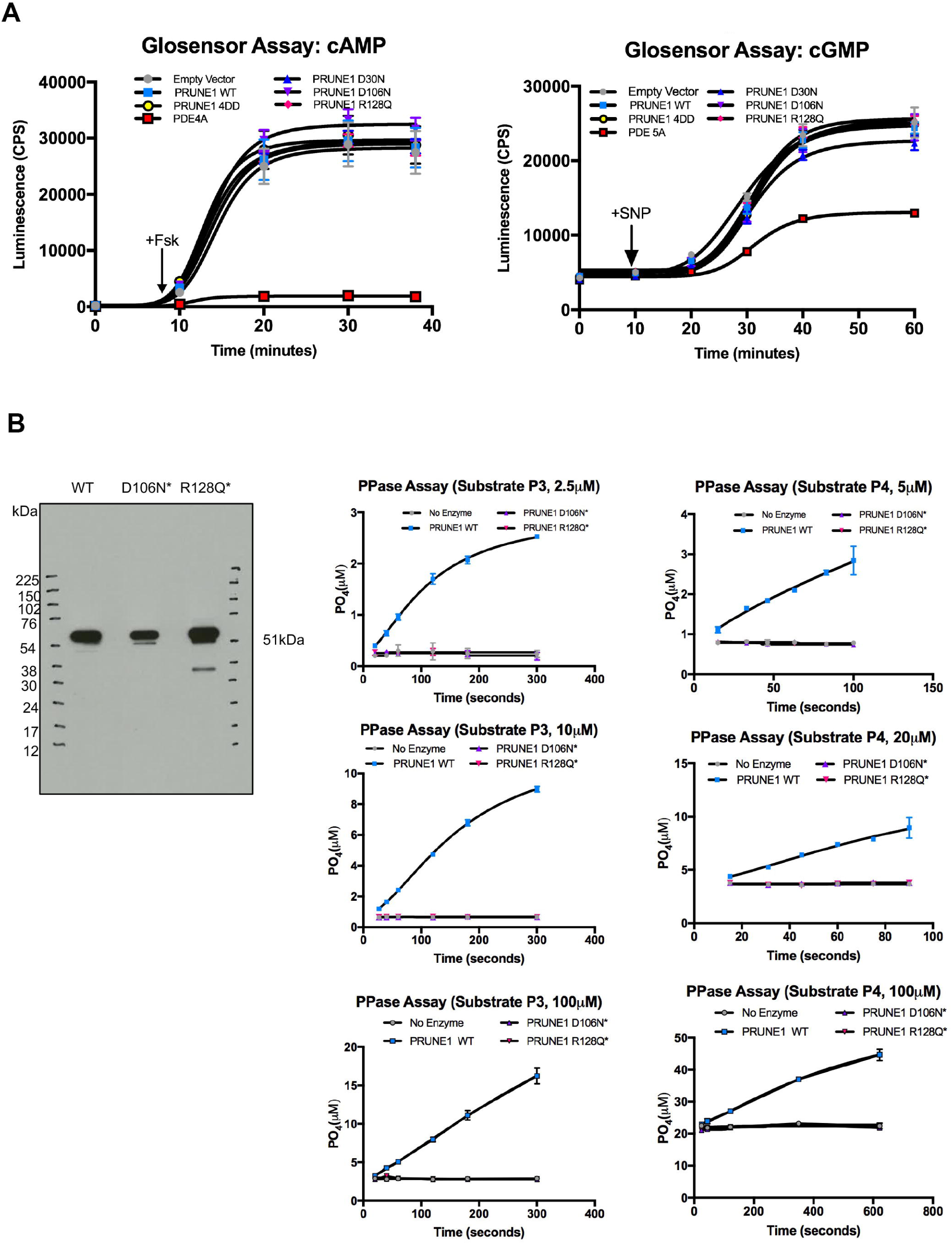
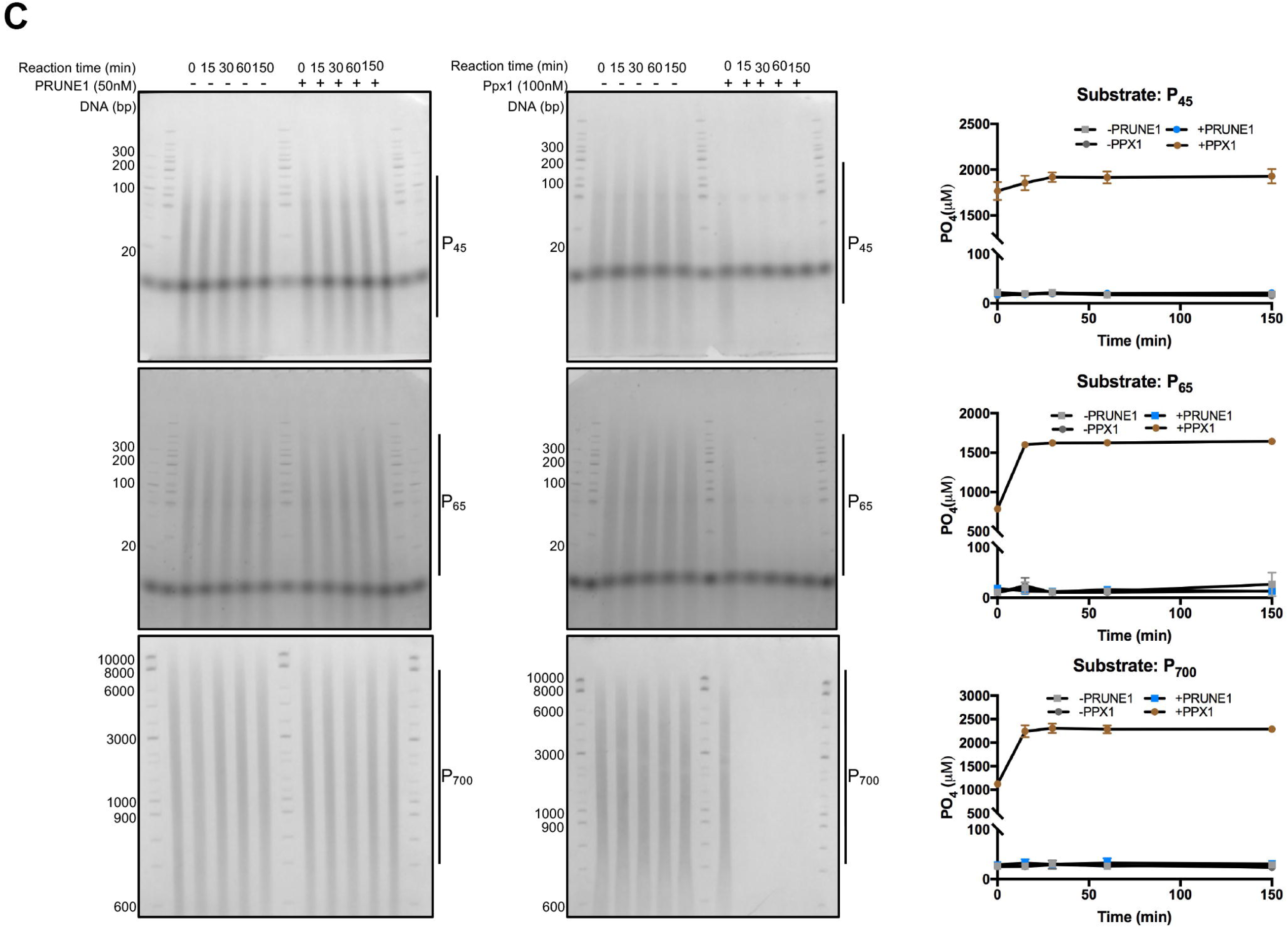
(**A**) HEK293 cells were transfected with pGlosensor-22F or pGlosensor-42F along with PRUNE1 wild type or mutant (D30N, D106N, R128Q, 4DD) constructs. PDE4A and PDE5A were used as positive controls for cAMP and cGMP assays respectively. Forskolin (Fsk, 10µM) or sodium nitroprusside dehydrate (SNP, 50µM) stimulated luminescence is plotted over time. (**B**) Immunoblotting of Expi-293 derived wild type PRUNE1 and *E*.*Coli* derived D106N* and R128Q* muteins (left panel). Short-chain exopolyphosphatase activity of wild type, D106N* and R128Q* mutants on P3 and P4 determined using fixed-time BIOMOL Green phosphate detection assay (middle and right panel). (**C**) Long-chain exopolyphosphatase activity detected by PAGE analyses. Synthetic polyphosphates at a final concentration of 2mM (with an average chain length of 45, 65 or 700) were treated with or without PPX1(100nM) or wild type PRUNE1(50nM). Reactions terminated at various time points (15, 30, 60, 150min) were resolved using 20% (for P_45_ and P_65_ reactions) or 6% (for P_700_) TBE gels (left panel). End-point reactions (terminated at 150min) were quantified using BIOMOL Green phosphate detection assay (right panel). Quantified data for Glosensor assay (panel A) and PPase assay (Panel B) represent mean ± SEM over 3 independent experiments with 6 technical replicates per sample.

**Figure S6.**
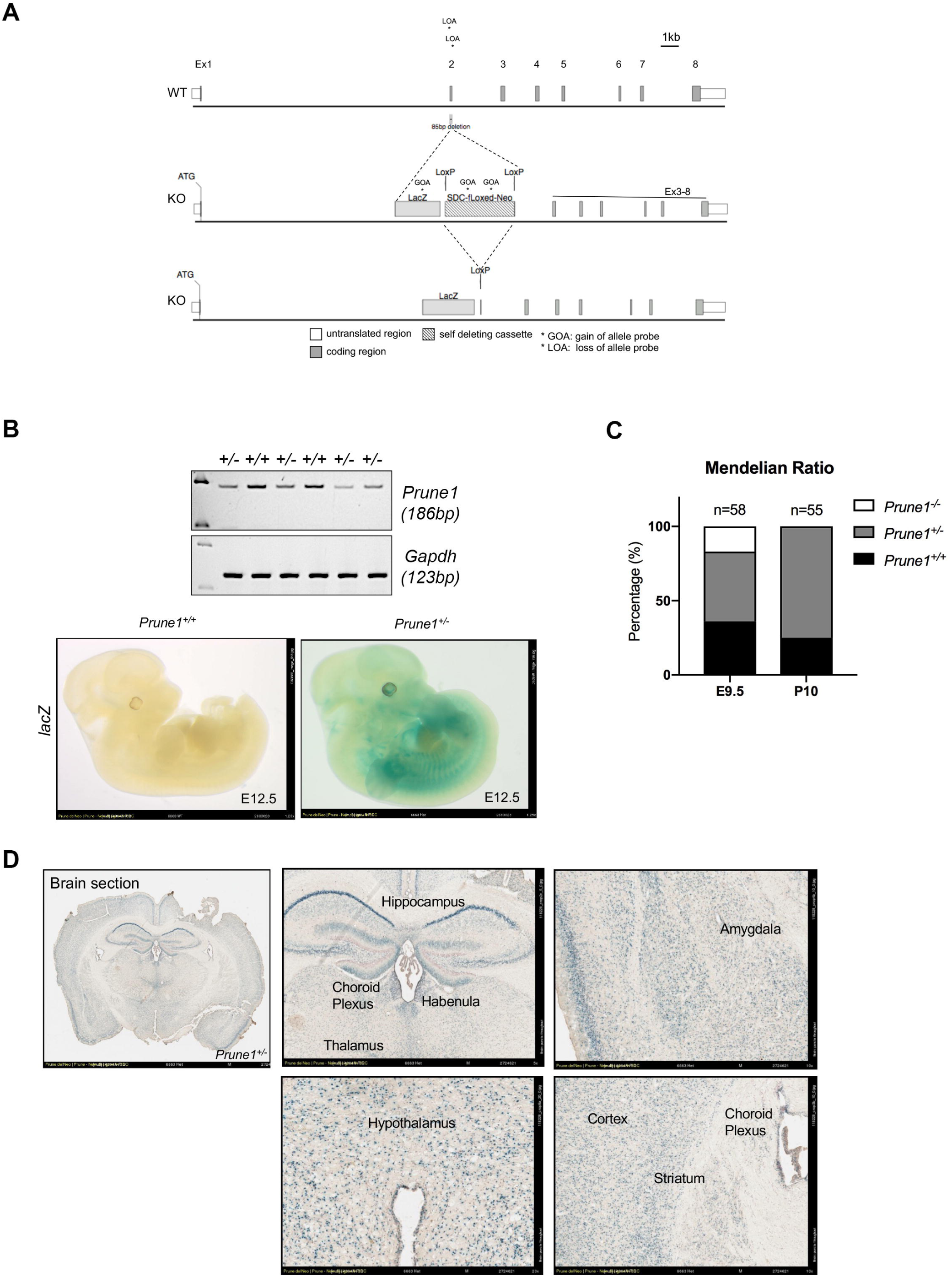
(**A**) Schematic showing homologous recombination mediated knockout strategy to generate *Prune1* null allele. **(B)** Developmental *Prune1* expression visualized by whole mount LacZ reporter staining in E12.5 *Prune1*^*+/-*^ embryos. Images are representative of multiple wild type and heterozygote littermates (n>5). **(C)** Gel electrophoresis of RT-PCR products using *Prune1* and *Gapdh* primers from cDNA derived from E9.5 embryos. **(D)** LacZ expression profiling of adult Prune1^+/-^ mice demonstrating widespread LacZ^+^ cells in the hippocampus, cerebellum, amygdala, hypothalamus and the cortex.

**Figure S7.**
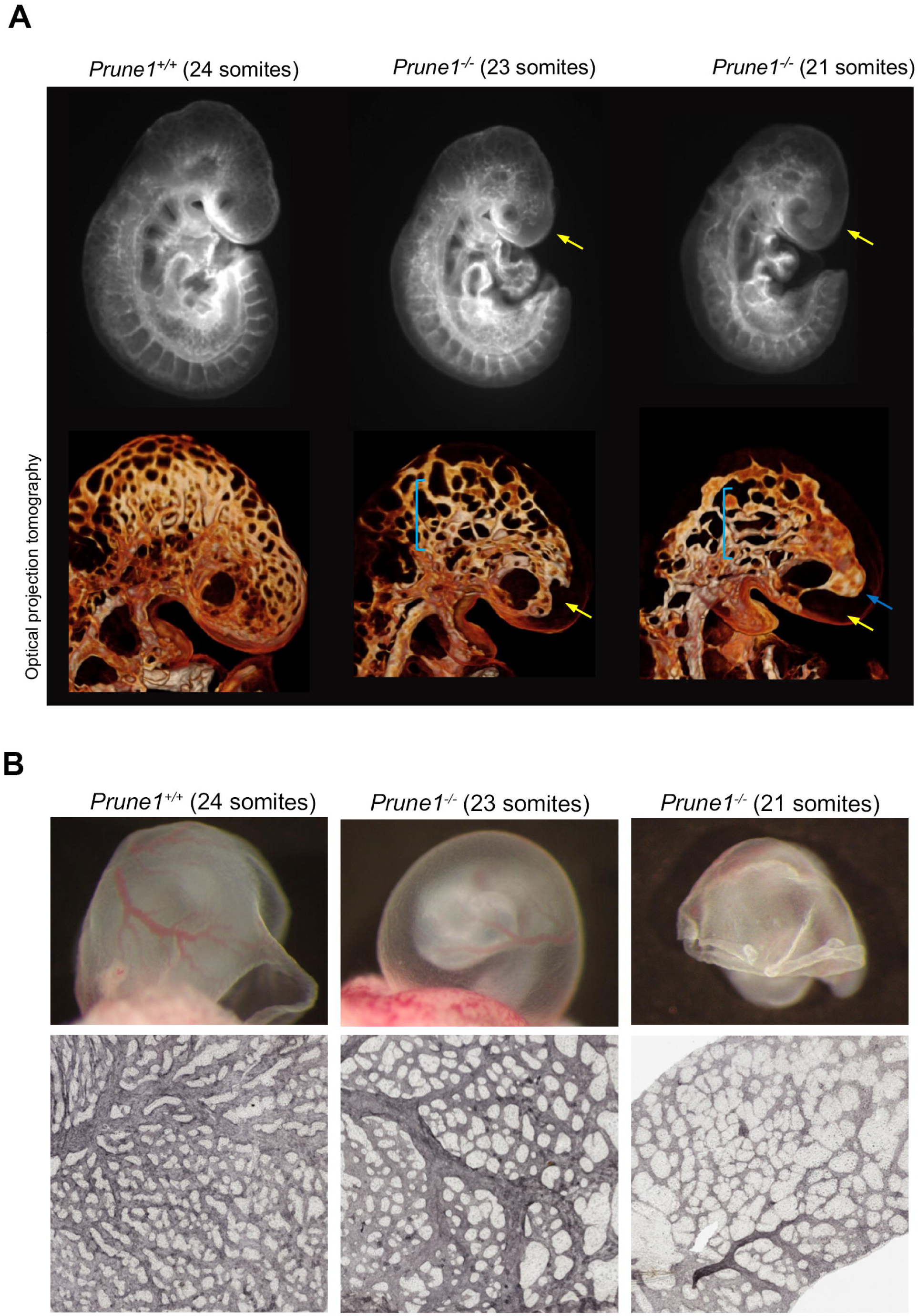
(**A**) Optical projection tomography revealed significant brain anomalies in *Prune1*^*-/-*^ embryos between E9-E10. Disrupted cephalic vascular plexus (yellow arrows) with large endothelial sheets (blue bracket) or sacs (blue arrows) observed in the *Prune1*^*-/-*^ embryos. (**B**) Brightfield images of freshly dissected yolk sacs from *Prune1*^*-/-*^ embryos between E9-E10 (top panel). Pecam1 staining further highlights defective vascular remodeling in the *Prune1*^*-/-*^ embryos as compared to wild-type. One *Prune1*^*+/+*^ and 2 *Prune1*^*-/-*^ embryos were analyzed for these experiments.

**Figure S8.**
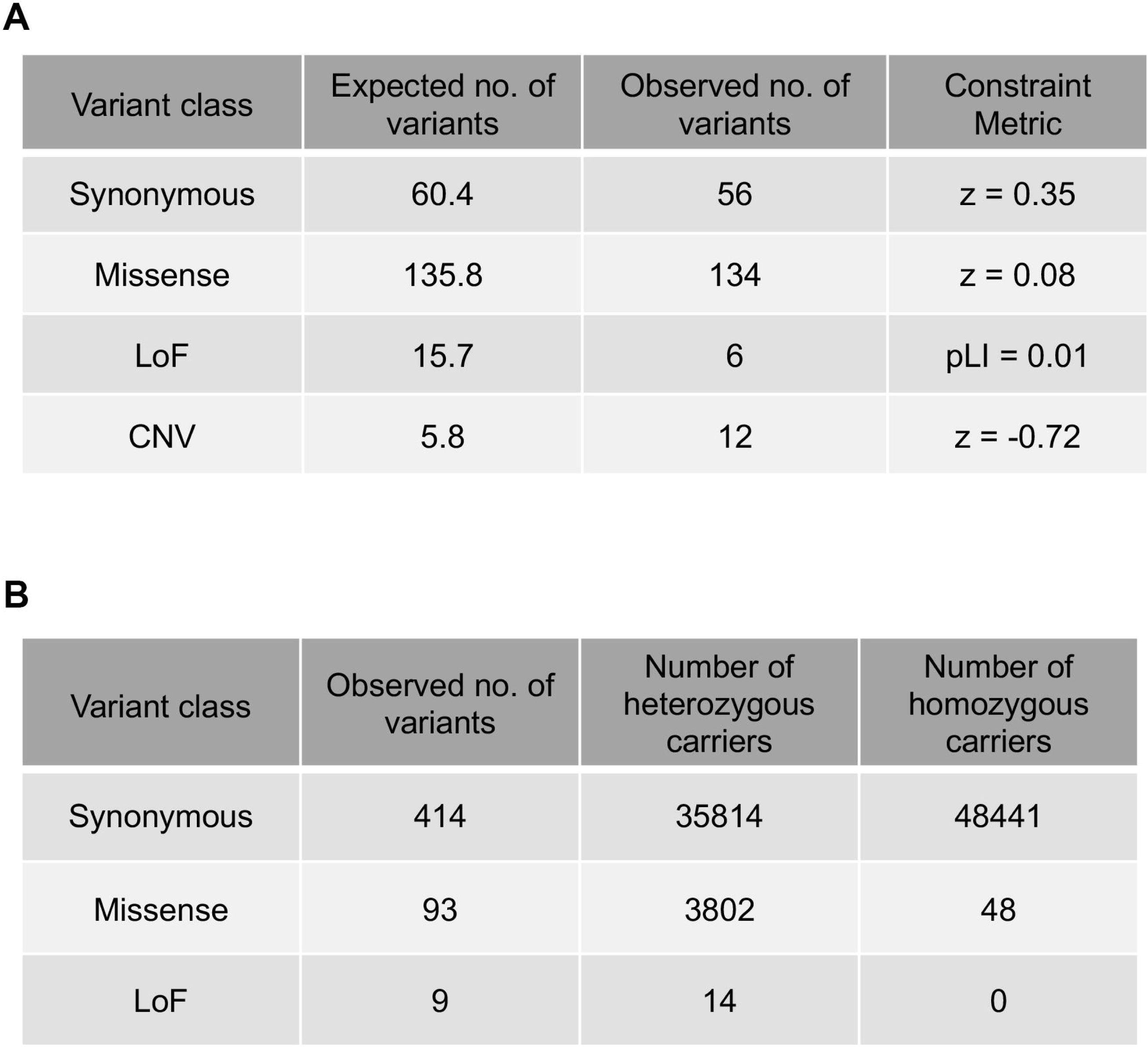
(**A**) Variant summary for *PRUNE1* showing expected and observed counts for different variant classes and gene-level constrain metrics for *PRUNE1* as reported by Lek *et al (28)*. ExAC constraint metric pLI = 0.01 for *PRUNE1* indicates that it is tolerant of heterozygous loss of function variation. (**B**) Analysis of protein-coding variation within *PRUNE1* as part of DiscovEHR study identified no homozygous carriers of LOF variants.

### Clinical description of probands in family SZ51

Patient 1 was born at 37 weeks after an uncomplicated pregnancy. Except for her occipitofrontal head circumference (11^th^ percentile), her height, weight, appearance, pulse, grimace, activity and APGAR scores (9 at 1min and 9 at 5min) were unremarkable at birth (Table. S2). Severe reflux due to a milk protein allergy was noted at 2 months. Developmental delay manifested by poor head control, not reaching for toys, and poor eye contact was observed by 3 months. At 4 months, she had an exaggerated startle, an aversion to various sensory stimuli, and no social smile. Furthermore, she developed upper extremity spasticity and intermittent esotropia, which became constant by 6 months requiring surgical intervention at 9 months. At 5 months, she was diagnosed with left hip dysplasia, which was treated at 10 months. By 6 months, she developed myoclonic seizures. EEG at the time revealed a few spikes, but no seizure activity. At 2.5 years, her myoclonic seizures worsened, and her EEG showed slow waking background and slow spike waves. Although her EEGs became progressively abnormal, the myoclonic seizures were better controlled with taking adrenocorticotropic hormone (ACTH) and vigabatrin (an inhibitor of gamma-aminobutyric acid aminotransferase). Developmentally, she could not sit independently, crawl, walk, or talk. She received a pureed diet, however due to poor weight gain, she was administered supplemental nutrition through a feeding tube. The patient was evaluated at the National Institutes of Health Undiagnosed Diseases Program (NIHUDP) at age 4 years. She presented with profound developmental delay, seizures, axial hypotonia and distal limb spasticity/hypertonia, severe kyphoscoliosis, exotropia, and severe gastroesophageal reflux disease (GERD) resulting in J-tube feeding. She had no purposeful movements and communicated with cries and grunts as her only vocalization. Serial MRI scans (at 4 months and 4 years) demonstrated progressive generalized cerebral and cerebellar atrophy (Fig. 1B). EEG showed infrequent right posterior temporal and left temporal spike waves and generalized background slowing. Measurement of mitochondrial respiratory chain enzyme activity revealed normal results once corrected for low citrate synthase activity (Table S3).

Patient 2 is the younger brother of patient 1 and has a dizygous twin sister who is unaffected (Fig. 1A). Conceived through *in vitro* fertilization, he was born at 38 weeks by elective C-section from an uncomplicated pregnancy. He was noted to be jittery at birth and also had marked stiffness of his feet. Like his older sister, he was found to have milk protein allergy and reflux by 2 months (GERD diagnosed at 4 months) as well as congenital intermittent esotropia. At 3 months, he appeared hyper-alert with eye popping when startled and exhibited intermittent down gaze associated with upper eyelid retraction. He was tremulous and had increased limb tone, brisk tendon reflexes (+3), bilateral Babinski signs and sustained ankle clonus. He began having seizures at 6 months of age. EEG showed a pattern of modified hypsarrhythmia with periodic synchronous discharges alternating with intervals bilaterally independent multifocal epileptiform discharges, background slowing, clusters of electro-decremental activity and loss of expected background organization during sleep and wakefulness (Table S2). He was treated with ACTH resulting in modest improvement of the seizures but did not obtain complete seizure control. Developmentally, he had severe delays including poor head control and the inability to sit independently or stand. Evaluation at the NIH at 20 months old evidenced developmental delay, seizures, mild intermittent exotropia, and kyphoscoliosis. He had poor head control, could not sit or stand, had no purposeful movements, communicated with grunts and cries, and was sensitive to unexpected or quick movements. MRI imaging showed cerebellar vermian atrophy at this age (Fig. 1B).

### Plasmids

PRUNE1 overexpression constructs were derived from full length human cDNA clone (Origene RC206865, NM_021222). D30N, D016N, R128Q and G174* mutations were introduced using the Agilent QuikChange II XL site-directed mutagenesis kit, according to the manufacturer’s protocol using primers listed in supplementary table 1. Resulting sequence verified constructs were subcloned into a CMV-driven mammalian expression vector (Origene PS100013) and used for overexpression studies. Similarly, PDE4A and PDE5A overexpression constructs were derived by subcloning wild type cDNA (NM_001111307.1 and NM_001083 respectively) into the PS100013 backbone. We generated PRUNE1 4DD construct as described by D’Angelo et al. (D’Angelo A, 2004) by subcloning a g-block harboring D28A, D106A, DHRP126-129AAAA, D178A into the PS100013 vector. Additionally, PS100013 vector without an insert, was used as an empty-vector control where indicated. GloSensor reporter constructs (pGloSensor-22F and pGloSensor-42F, respectively) were purchased from Promega.

### Recombinant protein expression

Wild type, D106N and R128Q muteins were produced by overexpressing C-terminal His10-tagged PRUNE1 constructs in Expi293 cells followed by purification via affinity chromatography. Briefly, constructs encoding PRUNE1 isoforms (1mg) were transfected into Expi293 cells (3 × 10^9^) using ExpiFectamine 293 transfection reagent (Gibco A14524) in a total culture volume of 1L. Cells were cultured for 4 days following transfection in Expi293 expression medium (Gibco A14351), pelleted by centrifugation (500g for 10 minutes) at 4°C. Cell pellet was resuspended in lysis buffer (25mM Tris-HCl, 500mM NaCl, pH 7.4) and lysed by sonication. Protein was purified using a 5mL GE HisTrap column and eluted with a linear gradient of 20-500mM imidazole in 20mM Tris-HCl, 150mM NaCl, 2mM DTT, pH 7.8. Fractions for use were chosen based on purity, as determined by Coomassie Blue staining. Selected fractions were further dialyzed for 16 hours against 20mM Tris/HCl, 50mM NaCl, 2mM DTT, pH 7.8 and quantified using absorption at 280nm. Purity was further confirmed by SDS-PAGE and Native PAGE. For SDS-PAGE, 0.5ug, 1.0ug, and 2.0ug of each fraction was resolved using Novex 4-20% Tris-Glycine precast gel (Invitrogen XP04205BOX) in a denaturing/reducing sample buffer and run to completion using a Tris-Glycine SDS running buffer. The gel was then stained using Bio-Safe Coomassie stain (Bio-Rad 1610786) and imaged. The procedure for Native-PAGE was identical, with the exception of a non-denaturing, non-reducing sample buffer being used, as well as the use of a gel running buffer that did not contain SDS.

D106N and R128Q muteins were successfully purified from overexpression in *E. coli*. Expression of wild type PRUNE1 was cytotoxic. Therefore, short chain exopolyphosphatase activity of the E. coli derived D106N and R128Q muteins was compared to the Expi293 derived wild type PRUNE1. Expression constructs encoding N-terminal His-tagged PRUNE1 isoforms under the control of IPTG-inducible T5-promoter were transformed into Rosetta 2 competent cells (Novagen 71397). Upon selection, cultures were grown until they reached an OD of 0.6. At this time, cells were induced with 1mM IPTG and grown overnight at 25°C and pelleted. To purify the D106N mutant, pelleted cells (1g) were resuspended in lysis buffer (50mM Tris-HCl, 500mM NaCl, pH 7.8) containing Roche Complete Protease Inhibitor Cocktail, benzonase, and PMSF. Cells were sonicated and centrifuged at 14,000 RPM for 30 minutes at 4°C to separate soluble lysate. 10ml of the soluble lysate was added to 0.5mL of equilibrated TALON metal affinity resin (Clontech 635502), rocked at 4° Celsius for 1 hour, loaded in a 10mL column, and washed three times with the lysis buffer. Elutions were then performed using 150mM imidazole in lysis buffer. *E*.*Coli* expressed D106N and R128Q and Expi293 expressed wild type PRUNE1, were then dialyzed for 20 hours against 20mM Tris/HCl, pH 7.8, 50mM NaCl, 2mM DTT using D-Tube Dialyzer tubes (Millipore 71507). The purified protein fractions were then quantified by BCA. Purity of resulting recombinant proteins were evaluated by Coomassie Blue staining or immunoblotting as described previously.

---

**Table S1.**
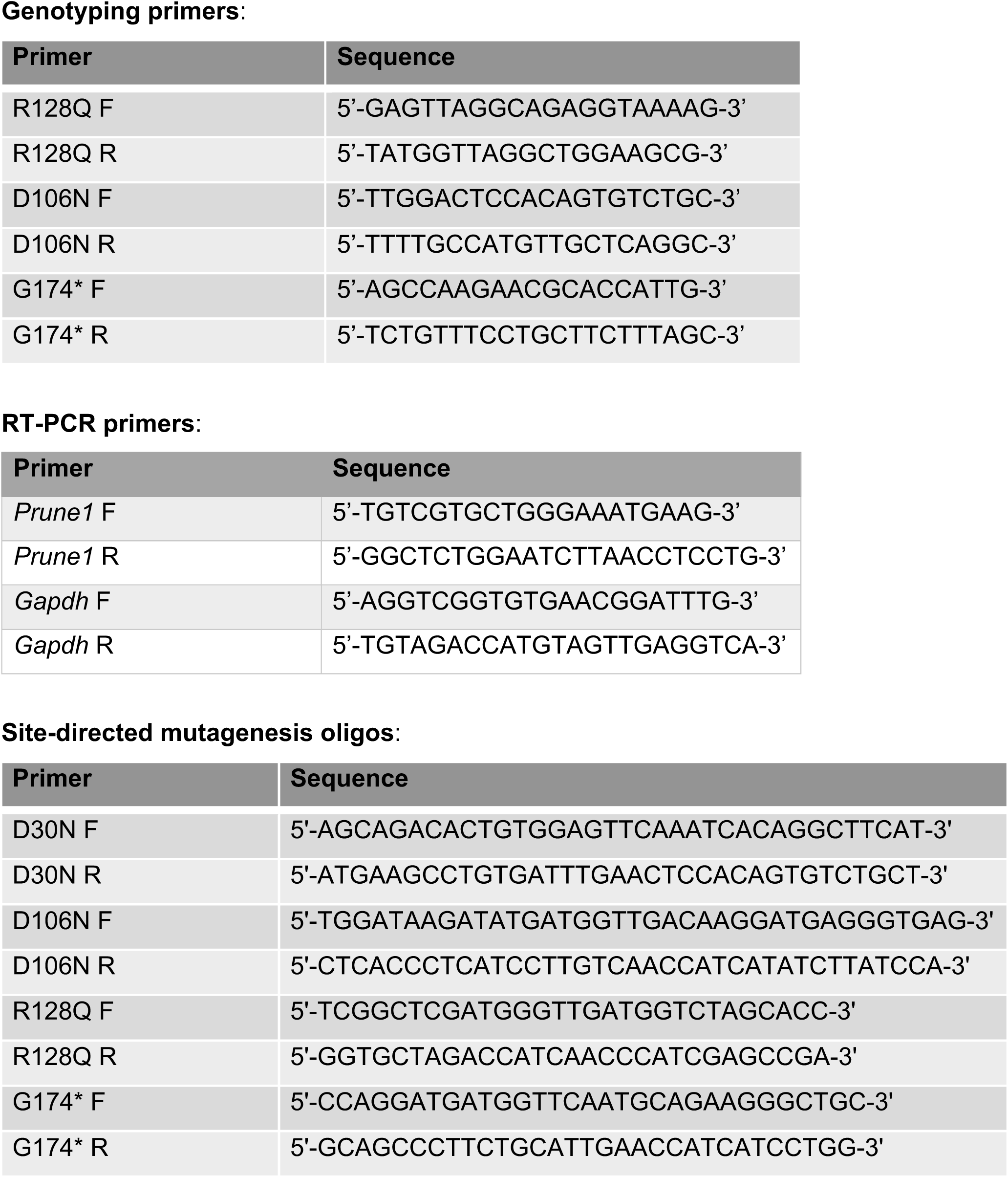

**G-block used for PRUNE1 4DD construct**: CAGATTACGCTGCGATCGCCATGGAGGACTACCTGCAGGGTTGTCGAGCTGCTCTGCAGGAGTCCCGACCTCTACATGTTGTGCTGGGAAATGAAGCCTGTGCTTTGGACTCCACAGTGTCTGCTCTTGCCCTGGCTTTTTACCTAGCAAAGACAACTGAGGCTGAGGAAGTCTTTGTGCCAGTTTTAAATATAAAACGTTCTGAACTACCTCTGCGAGGTGACATTGTCTTCTTTCTTCAGAAGGTTCATATTCCAGAGAGTATCTTGATTTTTCGGGATGAGATTGACCTCCATGCATTATACCAGGCTGGCCAACTCACCCTCATCCTTGTCGCCCATCATATCTTATCCAAAAGTGACACAGCCCTAGAGGAGGCAGTAGCAGAGGTGCTAGCTGCTGCTGCTATCGAGCCGAAACACTGCCCTCCCTGCCATGTTTCAGTTGAGCTGGTGGGGTCCTGTGCTACCCTGGTGACCGAGAGAATCCTGCAGGGGGCACCAGAGATCTTGGACAGGCAAACTGCAGCCCTTCTGCATGGAACCATCATCCTGGCCTGTGTCAACATGGACCTTAAAATTGGAAAGGCAACCCCAAAGGACAGCAAATATGTGGAGAAACTAGAGGCCCTTTTCCCAGACCTACCCAAGAGAAATGATATATTTGATTCCCTACAAAAGGCAAAGTTTGATGTATCAGGACTGACCACTGAGCAGATGCTGAGAAAAGACCAGAAGACTATCTATAGACAAGGCGTCAAGGTGGCCATTAGTGCAATATATATGGATTTGGAGGCCTTTCTGCAGAGGTCTAACCTCCTTGCAGATCTCCATGCTTTCTGCCAGGCTCACAGCTATGATGTCCTGGTTGCCATGACTATCTTTTTCAACACTCACAATGAGCCAGTGCGGCAGTTGGCTATTTTCTGTCCCCATGTGGCACTCCAAACAACGATCTGTGAAGTCCTGGAACGCTCCCACTCTCCACCCCTGAAGCTGACCCCTGCCTCAAGTACCCACCCTAACCTCCATGCCTATCTTCAAGGCAACACCCAGGTCTCTCGAAAGAAACTTCTGCCCCTGCTCCAGGAAGCCCTGTCAGCATATTTTGACTCCATGAAGATCCCTTCAGGACAGCCTGAGACAGCAGATGTGTCCAGGGAGCAAGTGGACAAGGAATTGGACAGGGCAAGTAACTCCCTGATTTCTGGACTGAGTCAAGATGAGGAGGACCCTCCGCTGCCCCCGACGCCCATGAACAGCTTGGTGGATGAGTGCCCTCTAGATCAGGGGCTGCCTAAACTCTCTGCTGAGGCCGTCTTCGAGAAGTGCAGTCAGATCTCACTGTCACAGTCTACCACAGCCTCCCTGTCCAAGAAGACGCGTTAAGCGGCCGC

**Table S2.** Clinical findings in patient 1 and patient 2. (OFC: occipitofrontal circumference).

|  | Patient 1 | Patient 2 |
| --- | --- | --- |
| Prune mutation Allele 1 | c.383G>A (p.Arg128Gln) | c.383G>A (p.Arg128Gln) |
| Prune mutation Allele 2 | c.G520T (p.Gly174*) | c.G520T (p.Gly174*) |
| Gender | Female | Male |
| Ethnicity | Caucasian | Caucasian |
| Consanguinity | No | No |
| Developmental Delay | Profound | Profound |
| Scoliosis/Kyphosis | Yes | Yes |
| Vision | Optic atrophy, esotropia | Mild intermittent esotropia |
| Muscle tone | Axial hypotonia, distal limb hypertonia | Increased limb tone |
| GERD | Severe | Yes |
| Seizures | Gelastic, complex partial, infantile spasms | Infantile spasms |
| Seizure onset | 6 months | 6 months |
| EEG | Background slowing, Infrequent temporal spike waves | Modified-hypsarrhythmia, multifocal epileptiform discharges |
| MRI | Moderate/severe progressive global atrophy | Mildly prominent lateral ventricles and sulci, thinner splenium and corpus callosum |
| Measurements at birth | OFC: 33cm (11<sup>th</sup> centile)<br>Weight: 3.4kg (48<sup>th</sup> centile)<br>Height: 54.61 cm (97<sup>th</sup> centile) | Not available |
| Age at Consultation | 4+1/12 years | 1+8/12 years |
| Measurements at consultation | OFC: 41.5 cm (-5.3 s.d.)<br>Weight: 15.1kg (24<sup>th</sup> centile)<br>Length: 94 cm (-2.0 s.d.) | OFC: 41 cm (-5.3 s.d.)<br>Weight: 12.8 kg (70<sup>th</sup> centile)<br>Length: 86 cm (77<sup>th</sup> centile) |

**Table S3.** Mitochondrial respiratory chain enzyme activity levels measured in dermal fibroblasts from patient 1 compared to values from age-matched controls. All values reported in nmoles/min/mg of protein.

| Enzyme tested | Patient's results | Control Range |
| --- | --- | --- |
| NADH dehydrogenase | 859 | 830-1222 |
| NADH cyto c reductase |  |  |
| Total | 239.7 | 216-302 |
| Rotenone sensitive | 87.4 | 92-178 |
| Succinate dehydrogenase | 5.36 | 6-7.96 |
| Succinate cyto c reductase | 1.52 | 2.93-5.43 |
| Cytochrome c oxidase | 9.8 | 13-21.2 |
| Citrate synthase | 39 | 53.7-81.9 |

